# Two distinct durable human class-switched memory B cell populations are induced by vaccination and infection

**DOI:** 10.1101/2024.11.29.624972

**Authors:** Cory A. Perugino, Hang Liu, Jared Feldman, Jess Marbourg, Thomas V. Guy, Anson Hui, Nicole Ingram, Julian Liebaert, Neha Chaudhary, Weiyang Tao, Catherine Jacob-Dolan, Blake M. Hauser, Zayd Mian, Anusha Nathan, Zezhou Zhao, Clarety Kaseke, Rhoda Tano-Menka, Matthew A. Getz, Fernando Senjobe, Cristhian Berrios, Onosereme Ofoman, Zachary Manickas-Hill, Duane R. Wesemann, Jacob E. Lemieux, Marcia B. Goldberg, Kerstin Nündel, Ann Moormann, Ann Marshak-Rothstein, Regina C. Larocque, Edward T. Ryan, John A. Iafrate, Daniel Lingwood, Gaurav Gaiha, Richelle Charles, Alejandro B. Balazs, Aridaman Pandit, Vivek Naranbhai, Aaaron G. Schmidt, Shiv Pillai

## Abstract

Memory lymphocytes are durable cells that persist in the absence of antigen, but few human B cell subsets have been characterized in terms of durability. The relative durability of eight non-overlapping human B cell sub-populations covering 100% of all human class-switched B cells was interrogated. Only two long-lived B cell populations persisted in the relative absence of antigen. In addition to canonical germinal center-derived switched-memory B cells with an IgD^-^CD27^+^ CXCR5^+^ phenotype, a second, non-canonical, but distinct memory population of IgD^-^CD27^-^ CXCR5^+^ DN1 B cells was also durable, exhibited a unique *TP63*-linked transcriptional and anti-apoptotic signature, had low levels of somatic hypermutation, but was more clonally expanded than canonical switched-memory B cells. DN1 B cells likely evolved to preserve immunological breadth and may represent the human counterparts of rodent extrafollicular memory B cells that, unlike canonical memory B cells, can enter germinal centers and facilitate B cell and antibody evolution.

**Graphical Abstract:** 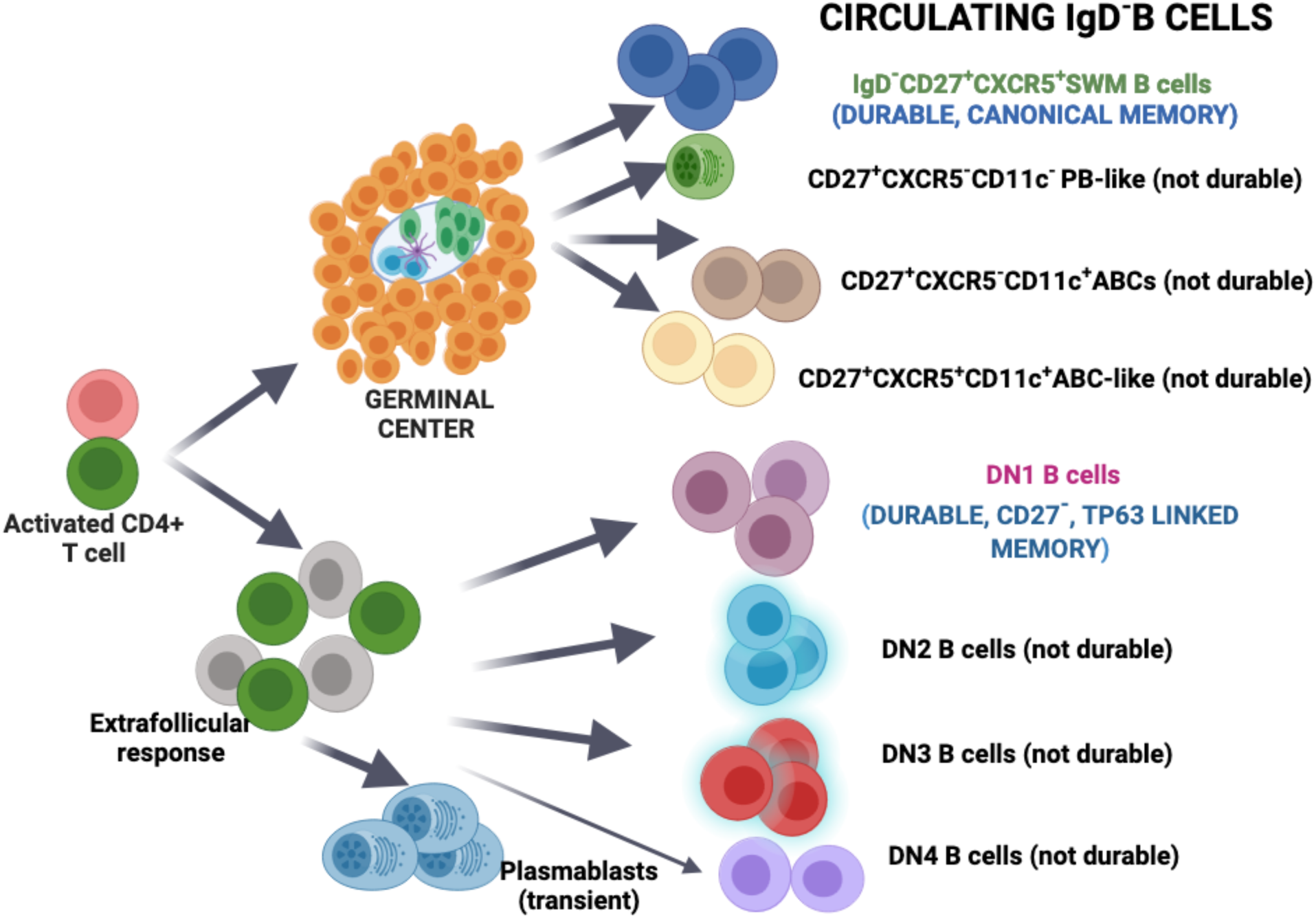

## Introduction

The traditional definition of a memory lymphocyte is of a durable antigen-specific cell that persists and can be activated when re-exposed to its cognate antigen (Ahmed and Gray, 1996). Canonical, durable memory B cells are traditionally thought to exhibit moderate to high levels of somatic hypermutation and to only emerge from a germinal center response.

There are many valid ways to characterize human memory B cells and numerous excellent reviews have described the properties of these cells (for example, Inoue and Kurosaki 2024, Weisel and Shlomchik, 2017, Cancro and Tomayko 2021). However, at this time, no categorization scheme for human memory B cells incorporates durability as a criterion. In humans, IgD^-^ CD27^+^ CXCR5^+^ CD21^+^ CD11c^-^ B cells are conventional switched memory (SWM) B cells believed to be derived from germinal centers based on the high levels of B cell receptor somatic hypermutation (SHM) that they exhibit relative to naïve B cells. These cells are relatively durable even in the context of subunit vaccination since durability cannot be convincingly established after live vaccines, infection, or autoimmunity, due to the persistence of antigen. Numerous studies, utilizing varying surface markers, have described many other different class-switched human B cell populations in the context of disease or infection many of these cells are referred to as memory B cells though persistence in the absence or relative absence of antigen has typically not been assessed.

There has been considerable interest in a heterogeneous atypical memory B cell subset often called Age-Related B cells or ABCs. In rodents, these were described as B220^+^ CD19^+^ B cells that do not express CD21, CD23, CD95 or CD43 (Hao et al. 2011), or as B220^+^ CD19^+^ CD21^-^ cells that express CD11c (Rubtsov et al., 2011). In the same study, Rubtsov and colleagues (2011) examined and defined human CD19^+^ CD21^-^ CD23^-^ CD11c^+^ IgD^-^ CD27^Hi^ human B cells as ABCs. The phenotype of ABCs is linked to the expression of the transcription factors T-bet and ZEB2 (Rubtsov et al., 2011, Cancro 2020, Dai et al., 2024). When ABCs are defined more loosely as Tbet-expressing B cells, they are heterogeneous (Cancro, 2020) and in humans these cells include not only IgD^-^ CD27^+^ CD11c^+^ cells, but also IgD^-^ CD27^-^ CD11c^+^ (i.e. double negative 2, DN2) and IgD^+^ CD27^-^ CD11c^+^ (activated naïve, aN) B cells and they are likely closely related to a fairly prominent population of cells that we will refer to below as ABC-like B cells.

When IgD^-^ CD27^+^ CD11c^+^ABCs were analyzed in a non-autoimmune and non-infectious context in influenza vaccination-based studies, they were characterized as germinal center-derived B cells since they exhibit robust SHM, but it was also suggested that these cells represent a transient activated B cell population based on their decline in the circulation about 8-10 weeks following vaccination (Lau et al., 2017). A second CD11c^+^ class switched B cell population in the IgD^-^ CD27^-^ pool has been studied especially in the context of systemic lupus erythematosus and these cells have been referred to as DN2 B cells (Kaminski et al., 2012; Jenks et al., 2018). Sanz and colleagues divided IgD^-^ CD27^-^ double negative (DN) B cells based on their expression of CXCR5 (or CD21, both used interchangeably) and CD11c into two prominent DN B cell subsets, namely, DN1 and DN2 B cells. DN2 B cells are IgD^-^ CD27^-^ CD11c^+^ CXCR5^-^ CD21^-^ B cells and similar to CD27^+^ ABCs, display clonal connectivity to plasmablasts. They have been shown to be expanded in many disease states. Based on these data, DN2 B cells represent a CD11c^+^ class switched B cell subset that can be distinguished from CD27^+^ABCs but likely represent plasmablast precursors that develop outside the germinal center based on their lower frequency of somatic hypermutation than SWM B cells (Jenks et al. 2018).

In more recent studies, we described four subsets of antigen-specific DN B cells in the context of SARS-CoV-2 infection, which extended the nomenclature to DN3 and DN4 subsets (Kaneko et al. 2020; Allard-Chamard et al., 2023). Sanz and colleagues have also described DN3 B cells in addition in the contexts of COVID-19 (Woodruff et al. 2020) and systemic lupus erythematosus (Jenks et al., 2021). DN3 B cells have a distinct plasmablast-like transcriptional profile and accumulate in non-lymphoid disease end organs far more readily than DN2 B cells (Allard-Chamard et al., 2023). DN1 B cells in this classification are IgD^-^ CD27^-^ CD11c^-^ CXCR5^+^ CD21^+^ B cells. Based on transcriptomic data demonstrating considerable similarity of DN1 B cells to IgD^-^ CD27^+^ CXCR5^+^ conventional switched memory B cells, these B cells were suggested to potentially represent a type of memory B cell population (Jenks et al., 2018). In more recent human studies using samples from convalescent COVID-19 patients (Zurbuchen et al., 2023), a relatively durable CD27^-^ memory B cell population was described (with a CD21^+^ CXCR5^+^ CD11c^-^ surface phenotype, consistent with DN1 B cells), supporting the notion that there may be an additional *durable* memory B cell population distinct from germinal center derived canonical IgD^-^ CD27^+^ CXCR5^+^ memory B cells. However, persisting antigen in any viral infection (or in autoimmunity) prevents firm conclusions regarding durability. In these previous studies, this subset of B cells was not characterized further in terms of identifying any unique transcriptomic features to convincingly distinguish them from canonical switched memory B cells.

By examining all eight class-switched B cell populations among antigen-specific CD19^+^ IgD^-^ B cells using CD27, CXCR5 and CD11c as markers (Table 1) in a previously uninfected vaccination cohort, as well as in acute and convalescent infection contexts, we were able to conclusively define only two relatively durable class-switched memory B cell populations and 6 short-lived B cell populations (not including plasmablasts). Among durable memory B cells, one type is the widely studied IgD^-^CD27^+^CXCR5^+^CD11c^-^ canonical class-switched memory B cell population that, as expected, has the hallmarks of being germinal center derived. The second durable class-switched memory B cell population, DN1 B cells, is likely derived outside the germinal center (possibly as part of an extrafollicular response) rather than in the course of a germinal center response. DN1 B cells have lower levels of replacement BCR mutations that span both CDR and FWR regions, they have longer CDRH3s than canonical SWM B cells, but they have clearly previously undergone clonal proliferation since they are more clonally expanded that conventional CD27^+^ memory B cells. These cells can also be clearly distinguished from canonical SWM B cells by a strong *TP63* transcriptome driven signature. These data define a second bona fide durable human class-switched memory B cell subset, that likely originates outside germinal centers and that may have evolved to contribute to immunological breadth. Six other human class-switched B cell subsets including the more widely studied ABC and DN2 B cells subsets, which can certainly be sustained by autoantigens and persistent infections, are nevertheless all intrinsically non-durable B cell subsets and perhaps should ideally be distinguished from durable memory B cell populations. The term “memory” is widely used in the B cell literature perhaps because issues of durability after antigen decline had not been previously addressed for most populations. In the context of class-switched B cells, we have generally attempted to restrict the use of this term in the remainder of this manuscript to the two durable B cell populations seen.

**TABLE 1.**
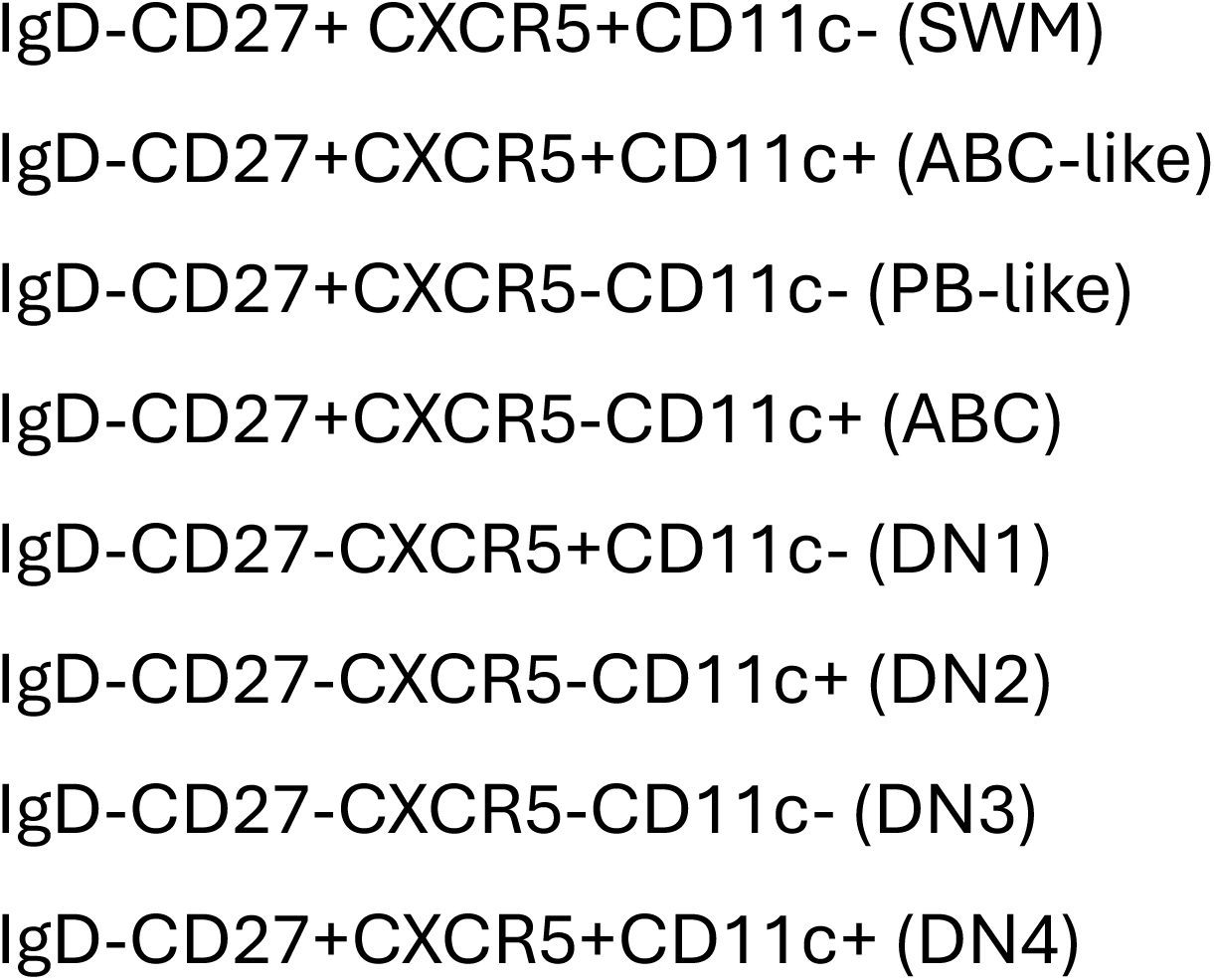
Class switched B cell subsets.

## Results

### Six distinct short-lived activated B cell populations are induced by mRNA vaccination and in acute viral infection

To understand the phenotypes, composition, and cellular kinetics of expansion and contraction versus persistence over time of antigen-induced human B cells, we interrogated SARS-CoV-2 spike-specific B cells from serially collected blood samples at baseline (pre-vaccination), soon after vaccination (1-2 weeks post-dose 1), at the time of vaccine boost (3-4 weeks post-dose 1), soon after vaccine boost (1-2 weeks post-dose 2), an intermediate duration after vaccine boost (3-4 weeks post-dose 2), and long-term after vaccine boost (6 months post-dose 2) from mRNA vaccine recipients (BNT162b2 or mRNA-1273) who had lacked SARS-CoV-2 nucleocapsid responses at baseline (i.e. SARS-CoV-2 naïve). These cohorts and time points were also studied alongside acute and convalescent COVID-19 patient samples to establish references for B cell subset composition in infection-induced systemic inflammation vs non-inflammatory contexts. Hierarchical gating of antigen-specific B cells was employed, and panel design was guided by the literature **(Supplemental Figure 1)**. Relatively few class-switched B cells were present within the spike-specific B cell gate at the pre-vaccination time point but those present were restricted to just two phenotypes: IgD^-^ CD27^+^ CXCR5^+^ (hereafter referred to canonical SWM) and IgD^-^ CD27^-^ CXCR5^+^ (double negative 1, hereafter referred to as DN1) **(Figures 1A-B)**.

**Figure 1.**
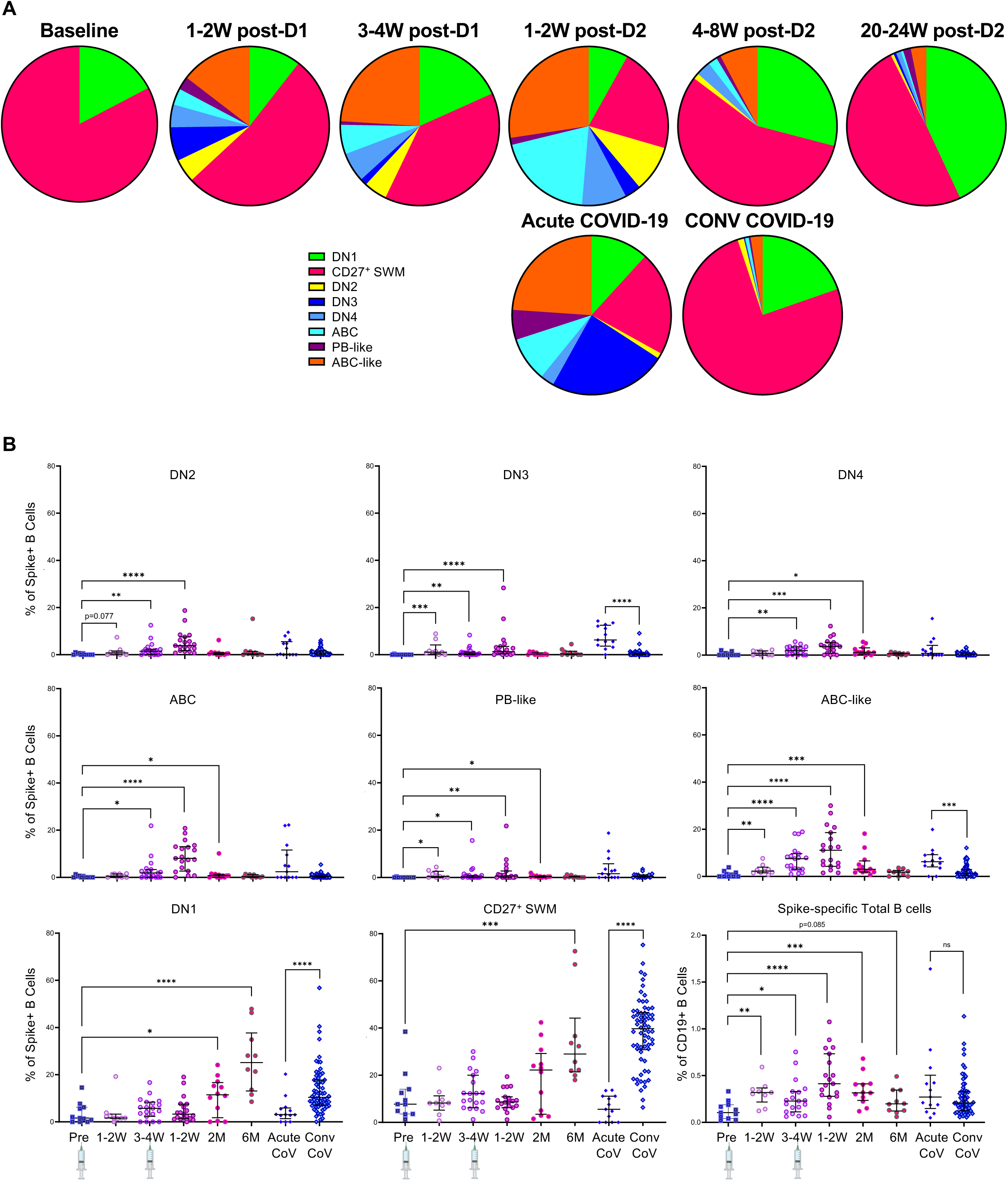
Two durable antigen-specific class switched B cell populations emerge after vaccination. A) Pie charts displaying spike-specific B cells profiled from the blood of previously uninfected vaccinees at baseline (n=11), 1-2 weeks post-dose 1 (n=9), time of dose 2 (n=19), 1 week post-dose 2 (n=19), 4-8 weeks post-dose 2 (n=12), and 24 weeks post-dose 2 (n=10), as well as from patients with acute (n=13) or convalescent from COVID-19 (n=66). The median of each B cell subset (as a proportion of all spike^+^ IgD^-^ B cells) from each time point are plotted as parts of a whole. Key: DN = double negative, IgD^-^ CD27^-^; ABC = age-associated B cells; CoV = COVID-19; Conv = convalescent. B) Scatter plots displaying the proportion of spike-specific B cell subset and total spike-specific B cells across the same cohorts depicted in part A. p-values were calculated using the Mann-Whitney test. Asterisks indicate p-values with values as follows: * 0.05 to 0.01; ** <0.01 to >0.001; *** <0.001 to >0.0001; and **** <0.0001.

When examining SARS-CoV-2 Spike antigen-specific B cells we observed six distinct short-lived antigen-specific B cell subsets (of which one is extremely minor) that appeared within 1-2 weeks following the first mRNA vaccination dose and generally receded to very low levels by 8 weeks after the 2^nd^ vaccination dose **(Figures 1A-B)**. These 6 short-lived B cell populations were defined by an activated phenotype based on varying combinations of the decreased expression of CXCR5 and CD21 (expression of these two antigens is generally coordinate following B cell activation) and/or the elevated expression of CD11c. For completeness and to ensure we accounted for 100% of class-switched spike-specific B cells, we included the very small population of CD27^+^ CXCR5^-^ CD11c^-^ B cells that have plasmablast-like features (most of these cells are frequently *either* CD20 low or CD38 high, referred to hereafter as “PB-like”), but this a very minor sub-population. Within the 3-4 weeks following the initiation of vaccination, DN3 B cells and PB-like B cells followed a time course similar to plasmablasts exhibiting rapid induction within 1-2 weeks followed by quick contraction by weeks 3-4. After the vaccination boost, the most prominent activated B cell subsets that appeared to most efficiently expand included the CD27^+^ CXCR5^-^ CD11c^+^ (age-associated B cells, hereafter referred to as ABCs), the closely related CD27^+^ CXCR5^+^ CD11c^+^ B cell population that we distinguish by flow but would often also be considered either directly as or as a variant form of CD27^+^ ABCs (we refer to these cells as “ABC-like”), CD27^-^ CXCR5^-^ CD11c^+^ (double negative 2, hereafter referred to as DN2), and to a lesser degree, CD27^-^ CXCR5^+^ CD11c^+^ (double negative 4, hereafter referred to as DN4) subset, suggesting these types of B cells are preferentially induced upon recall immune responses. The same 6 short-lived likely activated B cell sub-types were also observed in the setting of acute COVID-19 where the relative expansion of DN3 B cells was the most notable distinguishing feature compared to the mRNA vaccination cohorts, consistent with previous reports **(Figures 1A-B)**. The specific surface phenotypes and subset names where appropriate are detailed in **Table 1**.

### Two distinct durable switched-memory B cell populations are observed following vaccination and in the setting of acute infection and convalescence

By examining spike-specific B cells from SARS-CoV-2 naïve mRNA vaccine recipients 20-24 weeks after the 2^nd^ vaccine dose, we observed a stark paucity of all 6 short-lived B cell populations paired with the relative dominance of DN1 B cells and canonical CD27^+^ SWM B cells **(Figures 1A-B)**. Canonical CD27^+^ SWM B cells were more prominent, accounting for 20-80% of antigen-specific B cells at 20-24 weeks, whereas DN1 B cells accounted for 10-50%. Proportions of spike-specific B cells among total CD19^+^ B cells were higher following each vaccination dose **(Figure 1B)**, concordant with expansion of short-lived activated B cell subsets. In contrast, proportions of spike-specific B cells at the 6-month time point were similar to those of pre-vaccination donors. However, median proportions of DN1 B cells represented approximately 25% of all spike-specific B cells at 6 months compared to under 5% among pre-vaccination donors; those of canonical CD27^+^ SWM were approximately 30% vs 8% among pre-vaccination donors. As discussed above, canonical CD27^+^ SWM and DN1 B cells were the exclusive types of antigen-specific B cells present at the pre-vaccination time point. In contrast, when we examined receptor binding domain (RBD)-specific B cells, there were virtually no antigen-specific class-switched B cells in uninfected, unvaccinated individuals **(Figure S2A)**. These data suggest that all pre-existing class-switched memory B cells against the SARS-CoV-2 S2 domain, which has considerable homology to common cold coronavirus S2 domains, are dominated by canonical CD27^+^ SWM and DN1 B cells **(Figure S2B-C)**.

Both canonical CD27^+^ SWM and DN1 B cells appeared to reduce proportionally with the early emergence of short-lived activated B cell subsets (1-2 weeks after each vaccination dose) followed by their proportional increase seen at 4-8 weeks after vaccination boost. Canonical CD27^+^ SWM and DN1 B cells also dominated the antigen-specific B cell composition in the context of convalescence SARS-CoV-2 infection (Conv CoV) and were much less prominent in acute SARS-CoV-2 infection (Acute CoV). Consistent with previous reports, these data supported the existence of two types of memory B cells in humans that follow similar kinetic expansion after exposure to antigen, characterized by the surface phenotype IgD^-^ CXCR5^+^ CD21^+^ CD11c^-^ but distinguished by CD27 expression (Zurbuchen, 2023).

#### DN1 B cells represent a distinct relatively durable memory B cell population

We next studied a longitudinally collected cohort of convalescent COVID-19 patients with serially collected samples extending as far out as 258 days after symptom onset. Consistent with the findings from our vaccinee cohort, both canonical CD27^+^ SWM and DN1 B cells persisted over time with the former continuing to proportionally expand and the latter maintaining a plateau in most patients **(Figures 2A-B)**. We also examined circulating lymphocytes from a cohort of 24 healthy donors collected prior to the COVID-19 pandemic to understand the distribution of the six short-lived and two durable memory B cell populations in an antigen agnostic manner. Consistent with the notion of longevity of canonical CD27^+^ SWM and DN1 B cells, we found these two populations to dominate the circulating IgD^-^ B cell compartment with the former accounting for a median of 66% (IQR 56-76%) and the latter 18% (IQR 13-26%) of all IgD^-^ B cells **(Figure 2C-D)**. In contrast, each of the short-lived B cell types contributed median values of under 2% of all IgD^-^ B cells within this cohort, consistent either with their short-lived circulating lifespan or potentially their inability to re-circulate.

**Figure 2.**
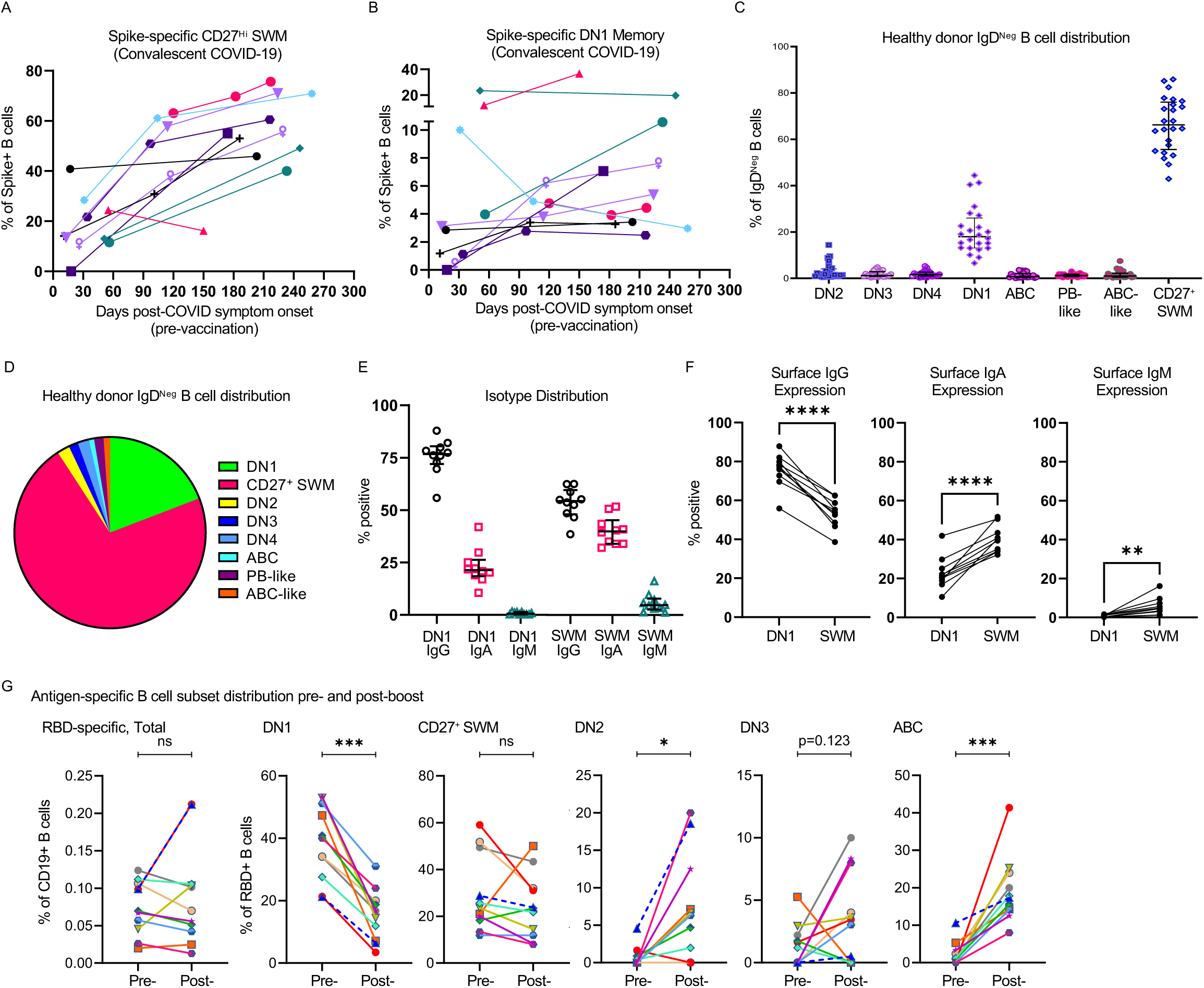
DN1 memory B cells and CD27^+^ SWM B cells accumulate at steady state and during convalescence. A-B) Dot plots with connecting lines displaying proportions of A) CD27^+^ SWM and B) DN1 B cells among all spike-specific B cells measured longitudinally across 11 patients during convalescence from COVID-19. Each color represents a different patient. C) Dot plot displaying proportions of all 8 non-overlapping, non-plasmablast, B cell populations among CD19^+^ IgD^-^ B cells across 24 pre-pandemic healthy donors. Lo/Lo and Hi/Hi refer to expression levels of CXCR5 and CD11c within the two CD27^+^ B cell subsets. D) Pie chart displaying the median values for each B cell subset as parts of a whole representing all non-plasmablast CD19^+^ IgD^-^ B cells profiled across 24 pre-pandemic healthy donors. E) Dot plot displaying the proportions of DN1 B cells expressing IgG, IgA, or IgM at the protein level among healthy donor PBMC samples (n=10). F) Dot plots with connecting lines comparing frequencies of IgG, IgA, and IgM surface expression between DN1 and SWM B cells. G) Dot plots with connecting lines showing pre- and post-boost proportions of RBD-specific B cells among all CD19^+^ B cells (first panel) or of each B cell subset among RBD-specific B cells (subsequent 5 panels) profiled across paired PBMC samples (n=11 from each time point). Each color represents a different donor. p-values were calculated using the Mann-Whitney test. NS = not significant. Asterisks indicate p-values with values as follows: * 0.05 to 0.01; ** <0.01 to >0.001; *** <0.001 to >0.0001; and **** <0.0001.

DN1 B cells are IgD^-^ by definition. To further understand if these cells are truly class-switched memory B cells, we characterized the isotype distribution of the surface BCR and found the majority to have class switched to IgG (mean 74%, S.D. 9.6%) and a minority to IgA (25%, S.D. 9.2%) with only a very small proportion to be unswitched with expression of surface IgM (1.3%, S.D. 0.86%) **(Figure 2E-F)**. Both canonical CD27^+^ SWM and DN1 B cells presumably rely on T cell help for their differentiation as supported by their class-switched phenotypes. We observed comparable high levels of surface CD40 expression in both memory subsets, supportive of their ability to interact with cognate T cells **(Figure S3)**. Both memory cell types also express high levels of the chemokine receptor CXCR5 and the selectin CD62L, similar to follicular naïve and unswitched memory B cells, suggesting tropism for secondary lymphoid organs **(Figure S3)**.

To better understand dynamic changes in the relative frequencies of these B cell memory types soon after antigen exposure, we examined the phenotype of antigen-specific B cells immediately before and after a 2^nd^ mRNA vaccination boost. All subjects were 6-7 months past their initial mRNA vaccination series with established B cell memory at the time of vaccine boost. In this relatively short interval, we did not observe significant expansion of antigen-specific B cells as a proportion of all CD19^+^ cells (only 3 of 11 donors showed clear proportional expansion within the entire B cell compartment). In contrast, we found marked shifts in the cellular composition of responding B cells despite the static overall fraction of antigen-specific B cells. DN1 B cells were consistently reduced and CXCR5^-^ CD11c^+^ short-lived activated B cells (ABCs and DN2 B cells) were consistently expanded across all 11 donors studied **(Figure 2G).** This observation suggests that DN1 B cells likely respond to antigen re-exposure by activation and differentiation and exhibit the transient dynamic decline expected of memory lymphocytes upon antigen re-exposure.

### DN1 B cells exhibit lower rates of somatic hypermutation and limited positive selection pressure than canonical SWM B cells

We wished to determine if the BCR sequence of DN1 compared to canonical SWM B cells could indicate their potential origins either from within germinal centers or from the extrafollicular response based on the degree of SHM these cells display.

We initially performed bulk IgH CDRH3 region BCR sequencing from FACS-purified naïve, DN1 and canonical SWM B cells using the Adaptive platform to quantitate the frequency of SHM. With these analyses, we found DN1 B cells to have a relatively low SHM frequency, closer to the low frequency seen in naïve follicular B cells and approximately 3-fold lower than the frequency found in canonical SWM B cells **(Figure 3A)**. In fact, the majority of DN1 B cells showed no mutations whatsoever within their CDRH3 regions **(Figure 3B)**. This was in stark contrast to canonical SWM B cells with nearly all B cells in these groups showing some degree of CDRH3 mutation and typically at a much higher frequency **(Figures 3A-B)**. The Adaptive platform provides sequence information on a limited portion of Ig VH genes - primarily the FWR3 and CDR3 regions - and the lengths of CDRH3 regions were seen to be longer in DN1 B cells (and similar to CDRH3 lengths in naïve follicular B cells) while CDRH3 length in SWM B cells was markedly reduced **(Figure 3C)** as has been reported before (Rosner et al., 2001; DeKosky et al., 2016).

**Fig 3.**
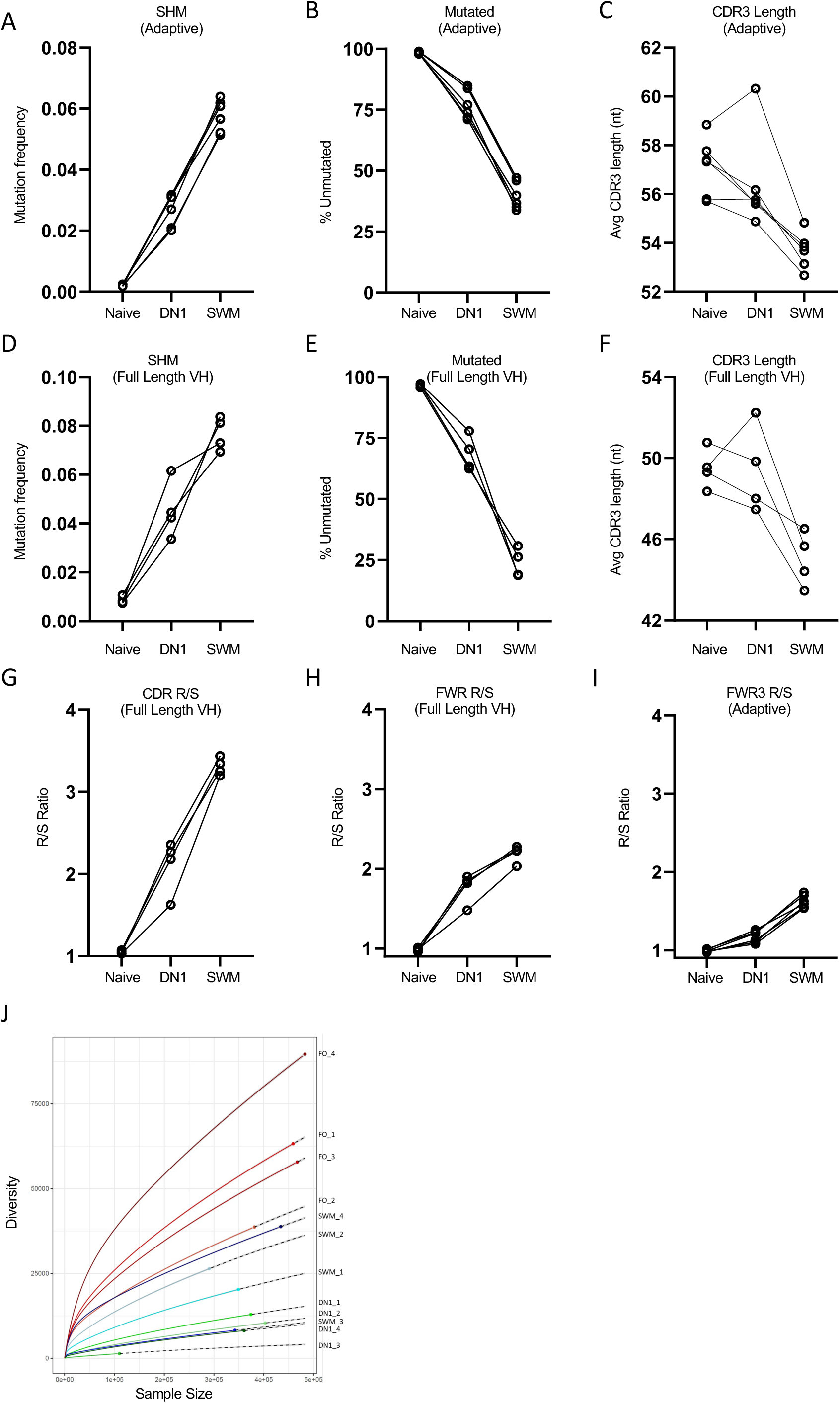
DN1 B cells exhibit lower SHM frequency, less positive selection pressure, and reduced repertoire diversity relative to canonical CD27+ SWM. A-C) Dot plots with connected lines displaying somatic hypermutation frequency, proportions of BCRs with unmutated germline sequences, and average CDR3 length among flow purified follicular naïve, DN1, and canonical SWM B cells across 6 healthy donor biologic replicates analyzed on the Adaptive platform. D-H) Dot plots with connected lines displaying (D) somatic hypermutation frequency, (E) proportions of BCRs with unmutated germline sequences, (F) average CDR3 length, (G) CDR replacement/silent (R/S) mutation ratio, and (H) FWR R/S mutation ratio among flow purified follicular naïve, DN1, and canonical SWM B cells across 4 different healthy donor biologic replicates analyzed by sequencing of the full length rearranged VH gene. I) FWR3 region R/S ratio displayed across the same biologic replicates in parts A-C using the Adaptive platform. J) Rarefaction curves of follicular, canonical SWM, and DN1 B cells displaying the BCR repertoire diversity (number of unique clones) relative to sequences analyzed for each population. Follicular naïve populations are depicted in tones of red, canonical SWM in tones of blue, and DN1 in tones of green. Each rarefaction curve is label with the cell population and associated biologic replicate the cells were sorted from. Key: CDR = complementarity determining region; nt = nucleotide; FWR = framework region; FO = follicular. Full length VH refers to sequencing of the complete VH gene (see Methods section).

To validate these findings and to be able to address issues related to selection of BCRs, we leveraged an orthogonal genomic DNA based full-length IgH V gene sequencing approach again using FACS-purified B cells from different healthy donors. When quantitating mutational frequency, a measurement of the extent of SHM, naïve B cells had the lowest averages of mutational frequencies as expected, DN1 B cells had an intermediate average, and switched memory B cells had the highest average, consistent with our initial findings **(Figure 3D)**. We observed a similar pattern when examining the proportion of non-mutated BCRs with follicular naïve B cells having the highest proportion, DN1 B cells showing an intermediate proportion, and canonical SWM having the lowest proportion of non-mutated BCRs **(Figure 3E)**. The longer average CDRH3 length of DN1 B cells compared to canonical SWM was also observed using this approach **(Figure 3F)**. Importantly, we also analyzed replacement to silent mutation ratios for complementarity determining regions (CDR) and framework regions (FWR). CDR R/S analyses revealed a degree of replacement mutation frequency among DN1 B cells that was intermediate between follicular naïve and canonical SWM B cells **(Figure 3G)**. FWRs in follicular naïve B cells exhibited a very low R/S ratio on average, while DN1 and canonical SWM B cells had higher R/S ratios with canonical memory B cells revealing slightly higher ratios on average relative to DN1 B cells **(Figure 3H-I).**

Examination of repertoire diversity found both DN1 and canonical SWM B cells to exhibit evidence of clonal expansion, as supported by reduced BCR diversity relative to follicular naïve B cells. Rarefaction curves were generated using a tool initially developed by ecologists to estimate the expected number of species from a sample of a larger diverse group (Sanders, 1968; Hurlbert, 1971). This approach can also help in the estimation of clonal diversity and clonal expansion for antigen receptor sequencing data. We created rarefaction curves for each sample of naïve follicular B cells, DN1 memory B cells or IgD-CD27+ switched memory B cells for which full length V gene sequences had been generated. We found that that DN1 memory B cells exhibited the highest amount of clonal expansion compared to switched memory B cells and follicular B cells **(Figure 3J)**. DN1 B cells consistently showed even lower diversity than canonical SWM B cells across the four biologic replicates studied.

### DN1 B cells exhibit a distinct TP63 linked transcriptional signature

While there are clear differences in rates of SHM between DN1 B cells and SWM B cells, we wished to ask if DN1 cells can, in a *robust* manner, be distinguished transcriptionally from canonical CD27+SWM B cells. Transcriptomes for both DN1 and SWM B cell populations have been compared in a limited manner so far (Jenks et al. 2018). Although we initially conducted CITE-seq and single cell transcriptomics on pre and post vaccination samples, given the broad similarities in the transcriptomes of the DN1 and CD27+IgD-switched memory B cell populations in the studies of Jenks et al. (2018) we recognized the need for deeper, more robust analyses. To determine whether a unique DN1 transcriptomic signature exists, we performed bulk RNA sequencing using sorted follicular naïve, DN1 and CD27^+^ SWM B cells isolated from six healthy control donor PBMCs. PCA clustering revealed strong separation between the three sorted cell populations, including previously undescribed separation between DN1 and canonical SWM B cells along PC2 (**Figure 4A**). This indicated the presence of a DN1 specific transcriptomic signature in this dataset. Comparing gene expression patterns between the three sorted B cell populations revealed thousands of differentially expressed genes (DEGs) between DN1 and naïve or canonical SWM B cells (**Figure 4B**). Of these DEGs, 1920 and 2421 genes were significantly upregulated and downregulated in DN1, respectively, compared to both naïve and SWM B cells (**Figure 4C**). Heatmaps of the topmost differentially expressed genes increased or decreased in DN1 are shown in **Figure 4D**. To better understand the unique features of DN1 B cells, we used Reactome Pathway Analysis (RPA) to identify which pathways were significantly enriched in the DEG sets consistently increased or decreased in DN1 (**Supplemental Figures 4A and B**). Notably, DEGs increased in DN1 were significantly enriched for pathways related to cell metabolism, including “the citric acid cycle and respiratory electron transport” and “cellular response to starvation.” DEGs decreased in DN1 were significantly enriched for pathways related to GTPase cycling.

**Figure 4.**
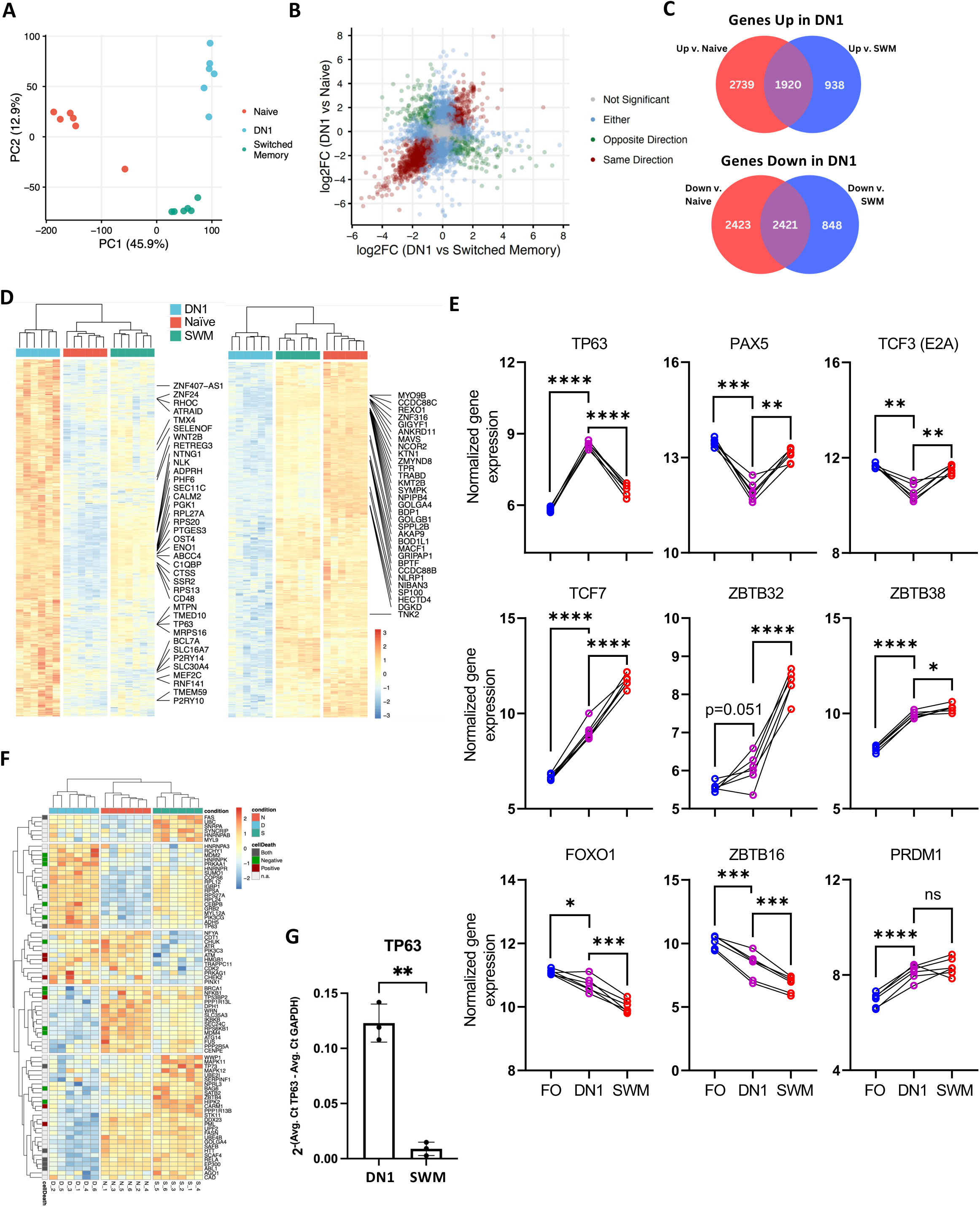
DN1 memory B cells are transcriptionally distinct from canonical SWM B cells and can be defined by a *TP63* linked signature. A) PCA plot of bulk transcriptomes shows strong separation with respect to sorted B cell subpopulations, including separation between switched memory (green) and DN1 (blue) B cells along PC2 (N=6/group). (B) Plot showing the number of differentially expressed genes (DEG) between DN1 and naïve (y-axis) or SWM B cells (x-axis), where each dot represents a gene. Colored dots represent genes with either the same (red) or opposite (green) direction of change in DN1 relative to both naïve and SWM. (C) Venn diagrams showing the number of significantly differentially expressed genes (DEG) that are increased (top) or decreased (bottom) in DN1 B cells compared to naïve and/or SWM B cells identified via bulk RNAseq. (D) Heatmaps showing the topmost DEG either increased (left) or decreased (right) in DN1 compared to the other B cell populations. (E) Plots showing VSD-normalized counts (log-scale) of gene regulators broadly associated with B cell differentiation comparing transcript expression levels across the three B cell populations. (F) Heatmap showing the expression of a TP63-dependent gene signature across the three B cell populations, where “cellDeath” indicates whether the indicated gene is found in the Gene Ontology (GO) annotation of negative (green) or positive (red) regulation of cell death, or both (gray). (G) Validation of *TP63* expression in DN1 and SWM B cells using qPCR.

To identify potential drivers of the DN1 transcriptomic signature, we used Regenrich to evaluate gene regulators significantly associated with the identified DEGs (Tao et al. 2022). Of the significantly associated gene regulators identified, only *TP63* was strongly increased specifically in DN1 whereas others such as TCF7 and ZBTB16 showed intermediate expression levels among DN1 in between follicular naïve and canonical SWM **(Figure 4E)**. *TP63* belongs to a family of transcription factors associated with regulation of cell proliferation and cell death known as the p53 family, which also includes *TP53* and *TP73*. Interestingly, *TP53* and *TP73* were not similarly increased in DN1 B cells (**Supplemental Figure 4C**). Because p63 signaling is associated with both positive and negative regulation of cell death, we explored what effect increased *TP63* expression might have on DN1 B cells by evaluating the expression of a *TP63*-driven gene signature across all three B cell populations. This TP63-dependent signature was obtained from the Harmonizome 3.0 database (Rouillard et. al, 2016). This identified sub-clusters of genes within the *TP63* signature that demonstrated increased or decreased expression specifically in DN1 (**Figure 4F**). Of the 19 genes specifically increased in DN1 B cells, six were present in the Gene Ontology (GO) annotation “negative regulation of cell death” while none were found in the GO annotation “positive regulation of cell death.” These 19 genes increased only in DN1 B cells included *PRKAA1*, the gene that encodes AMPK. Increased *PRKAA1* expression corresponded with a reduction in *MTOR* expression specifically in DN1 B cells **(Figure 4F)**, which relates to previous studies reporting that metabolic reprogramming of B cells, specifically AMPK activation, supports differentiation of unswitched memory B cells to DN B cells (Torigoe et al. 2017). TP63 has been linked to anti-apoptotic signaling in murine B cells and linked to the induction of BCL-2 (Lantner et al., 2007). The differential expression of *TP63* in DN1 B cells was confirmed using a quantitative RT-PCR approach **(Figure 4G)**. Overall, these data suggest that DN1 B cells possess a unique, *TP63*-driven gene signature that may contribute to their increased durability relative to other B cell subsets as previously described.

## Discussion

Our studies reveal that there are two durable class-switched human memory B cell populations that emerge after T-B collaboration, including one that is distinct from the canonical long-studied population that emerges from germinal centers. Not only is the data on durability supported by vaccination studies, it is also consistent with studies on infections and robustly supported by the examination of the accumulation, at steady state, in large numbers of apparently healthy individuals, of discernible class-switched B cells. We would suggest that in humans there are only two class switched B cell subsets that would fulfill the strict definition of memory by the criteria laid out by Ahmed and Gray (1996). Human IgD^-^ CD27^-^ CXCR5^+^ CD11c^-^DN1 B cells likely emerge outside germinal centers, as suggested by their relatively low frequency of SHM and low R/S ratios compared to canonical SWM B cells. These DN1 B cells are transcriptionally distinguished by the TP63 transcription factor which can be linked to specific downstream genes, including anti-apoptotic genes, that likely contribute to the distinct biology of DN1 memory B cells. Like canonical SWM B cells, DN1 memory B cells do not express TBX21 and ZEB2 which are, throughout lymphocyte biology, important markers of activated lymphocytes that reciprocally regulate TCF7 (Dominguez et al., 2015). It would be appropriate to recognize that the maintenance of intrinsically non-durable DN2 cells and CD27^+^ ABCs in disease contexts likely reflects the ongoing triggering of naïve or memory B cells by antigen and the repeated induction of ZEB2 and TBX21. Of course, in autoimmune subjects, the ubiquitous availability of antigen can lead to the persistence of intrinsically non-durable B cell subsets.

We suspect, as has been indicated in the literature, that DN1 B cells like DN2 (and DN3 B cells) emerge from the extrafollicular response (Jenks et al. 2018). Extrafollicular B cell responses in humans are harder to define because of the difficulty in micro-dissecting lymphoid organs for studies based on the geographical location of cells in secondary lymphoid organs. In our own multicolor immunofluorescence studies, albeit on fixed tissue sections and not performed using confocal imaging, IgD-CD27-BCl-6-B cells in secondary lymphoid organs were primarily localized outside germinal centers even in control subjects (Kaneko et al., 2020). Nevertheless, the usage here of the term “extrafollicular” should be considered operational rather than geographic. CD27 on human non-plasmablast, class-switched B cells may represent a surface marker for generally establishing the GC origin of canonical SWM B cells and some other activated B cell subsets. Average rates of SHM have therefore been used as a surrogate for GC or non-GC origin of activated and memory B cells.

Even with these caveats, we postulate an “extrafollicular” or possibly more narrowly, an “extra-germinal center” origin for human switched B cell subsets with some caution. Given the higher mutation frequency and replacement to silent mutation ratios of DN1 memory B cells compared to follicular naïve B cells, assuming mutations inherently have close to a 1 to 1 R/S ratio without selection, the results could suggest that DN1 B cells have undergone some degree of somatic hypermutation and selection, though not to the degree of canonical SWM B cells. This might theoretically have occurred either outside germinal centers or in germinal centers. In the latter scenario, after a few rounds of somatic hypermutation and selection, DN1 B cells could be postulated to have left the germinal center prematurely. However, a large number of DN1 B cells lack somatic mutations and we believe an origin in a distinct extra-germinal center milieu, likely the extrafollicular T-B boundary, may be permissive for TP63 induction and the development of durable DN1 memory B cells.

The studies of Shlomchik and colleagues on extrafollicular memory B cells in mice have revealed some crucial and important insights regarding the likely unique function of this category of memory B cells (Callahan et al., 2024). It should be noted that a large fraction of the murine extrafollicular memory B cells identified in those studies are IgM+ B cells. These studies reveal that extrafollicular memory B cells can enter germinal centers while GC-derived memory B cells cannot. The extrafollicular response may thus contribute crucially to durable B cell responses in many unique ways.

It has not been previously considered if there is any distinct, clearly distinguishable and durable class switched memory B cell subset with lower levels of somatic hypermutation and greater immune diversity that may exist potentially and primarily to maintain immunological breadth and to possibly feed into new cycles of germinal center responses with repeated exposure to antigens. We speculate that DN1 memory B cells have evolved to retain broad memory to inciting pathogens so that the same or variant pathogens can in subsequent infections utilize these cells to more robustly enter new germinal centers and to initiate affinity matured antibody responses. Such notions can best be tested in appropriate rodent models and indeed murine DN1 like B cells may be a subset of the cells studied by Callahan et al. (2024).

Overall, these data indicate that DN1 B cells are a durable class switched human memory B cell population that is likely derived through T-dependent B cell activation in the course of an extrafollicular response in secondary lymphoid organs. Although these cells exhibit limited SHM they undergo relatively large clonal expansions. DN1 B cells can be distinguished from canonical CD27^+^ SWM B cells since the latter exhibit high SHM, shorter CDRH3s on average, less B cell clonal expansion and have the hallmarks of being derived from germinal centers. Additionally, we establish here a TP63-linked transcriptome and associated anti-apoptotic signature that defines and distinguishes DN1 memory B cells from canonical SWM B cells. These data suggest an important evolutionary role for this memory B cell subset in maintaining broad immunologic antigen recognition breadth in contrast to the narrower affinity-matured recognition mediated by canonical SWM B cells.

The p63 protein family has been best studied in epithelial cells and most lymphocyte subsets do not express the *TP63* gene. Intriguingly, in many different categories of epithelial cells, differential promoter usage in the *TP63* gene can give rise to two major categories of p63 protein; one arising from longer transcripts that encodes TAp63, which possesses an N-terminal transactivation domain. This protein has an anti-apoptotic function. Usage of a distal promoter facilitates the generation of ΔNp63, a protein that mediates epithelial stemness in many different epithelial tissues (Senoo at al., 2007). It remains to be seen, potentially by the study of genetically engineered mice, whether *TP63* in the B lineage sustains a stem-like self-renewing memory B cell population.

## Acknowledgements

This work was supported by grants from the NIH, the HMS-Abbvie Consortium, the CDC (U01CK000490 and U01CK000633) the Lambertus Family Foundation and the Massachusetts Consortium for Pathogen Readiness.

## Author Contributions

Study Design/conceptualization: SP, CAP, JM; Execution: CAP, HL, JM, TVG, NI, JL, ZM; Reagent Generation: JF, CJ-D, BMH, AS, ABB, DL, AGS; Computational analyses, design and integration: AP, AH, NC, WT; Patient recruitment and sample processing: VN, RC, JAI, AM-R, AM, KN, MBB, JEL, DRW, ZM-H, OO, CB, FS, MAG, RT-M, CK, ZZ, AN. Writing, integration: SP, CAP, JM.

## Declaration of Interests

SP is on the Scientific Advisory Boards of Paratus Inc., BeBiopharma Inc, Octagon Therapeutics and AbPro Inc., all activities unrelated to the studies described here.

## STAR* Methods

### KEY RESOURCES TABLE

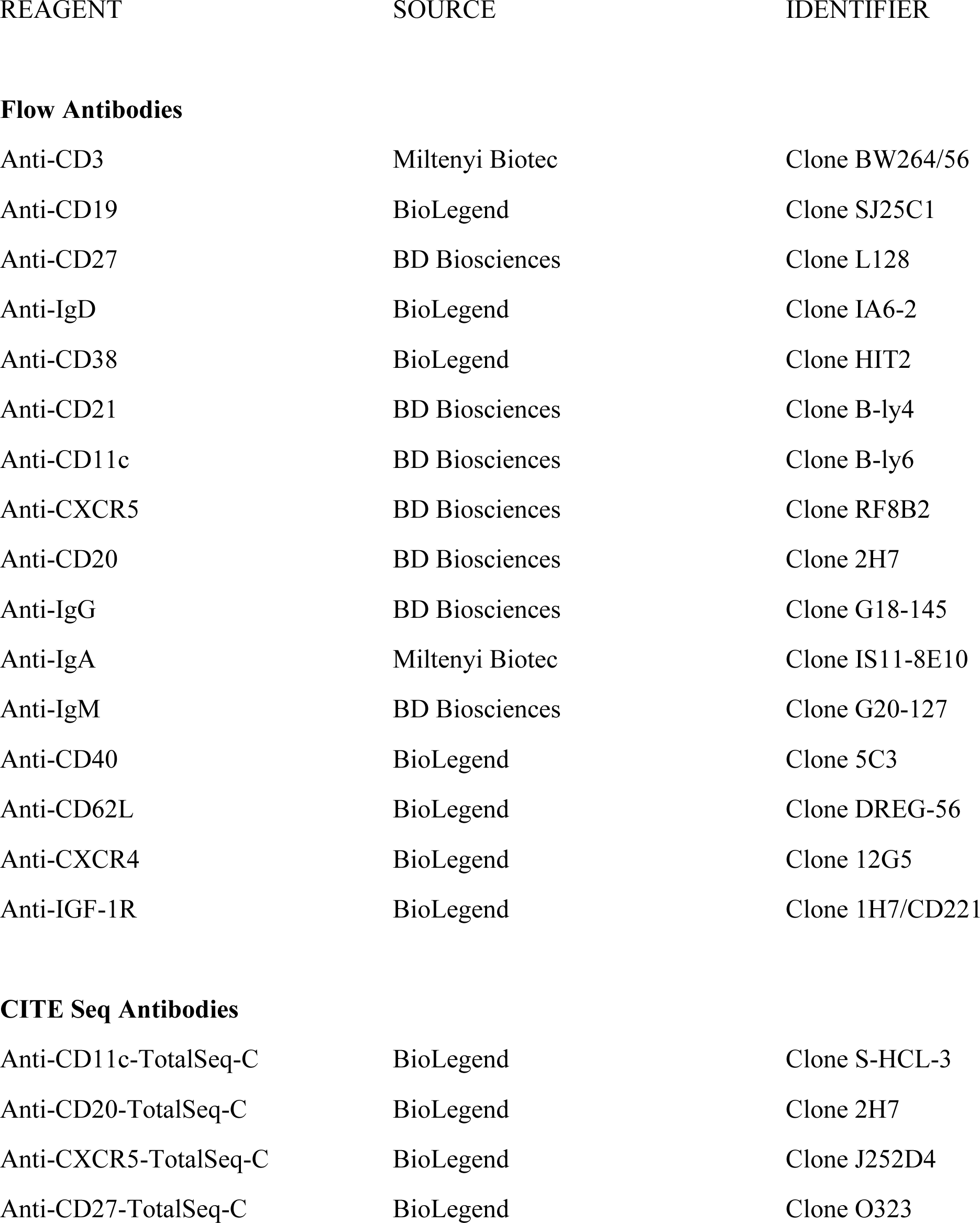
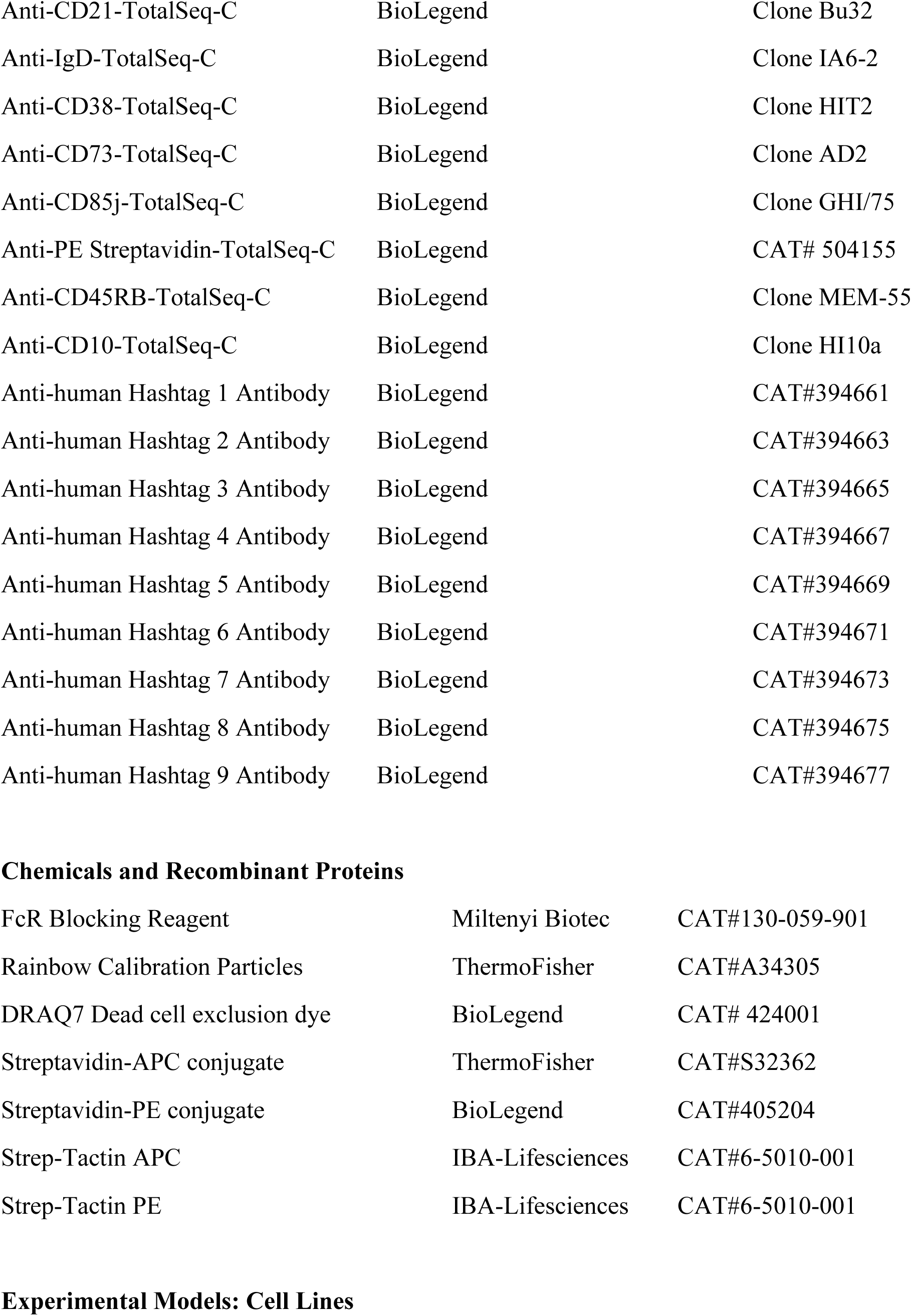
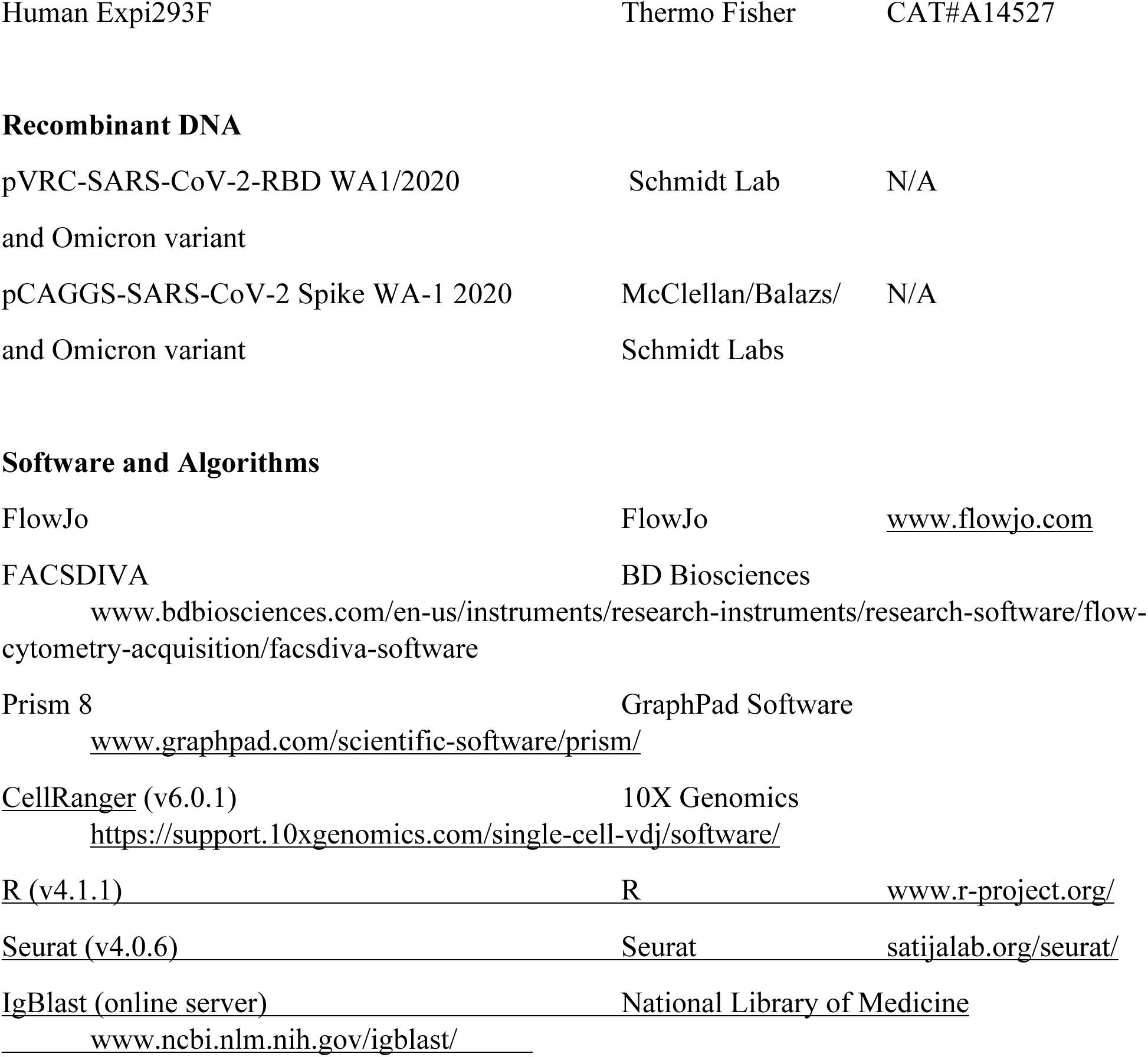

| REAGENT | SOURCE | IDENTIFIER |
| --- | --- | --- |
| **Flow Antibodies** |  |  |
| Anti-CD3 | Miltenyi Biotec | Clone BW264/56 |
| Anti-CD19 | BioLegend | Clone SJ25C1 |
| Anti-CD27 | BD Biosciences | Clone L128 |
| Anti-IgD | BioLegend | Clone IA6-2 |
| Anti-CD38 | BioLegend | Clone HIT2 |
| Anti-CD21 | BD Biosciences | Clone B-ly4 |
| Anti-CD11c | BD Biosciences | Clone B-ly6 |
| Anti-CXCR5 | BD Biosciences | Clone RF8B2 |
| Anti-CD20 | BD Biosciences | Clone 2H7 |
| Anti-IgG | BD Biosciences | Clone G18-145 |
| Anti-IgA | Miltenyi Biotec | Clone IS11-8E10 |
| Anti-IgM | BD Biosciences | Clone G20-127 |
| Anti-CD40 | BioLegend | Clone 5C3 |
| Anti-CD62L | BioLegend | Clone DREG-56 |
| Anti-CXCR4 | BioLegend | Clone 12G5 |
| Anti-IGF-1R | BioLegend | Clone 1H7/CD221 |
| **CITE Seq Antibodies** |  |  |
| Anti-CD11c-TotalSeq-C | BioLegend | Clone S-HCL-3 |
| Anti-CD20-TotalSeq-C | BioLegend | Clone 2H7 |
| Anti-CXCR5-TotalSeq-C | BioLegend | Clone J252D4 |
| Anti-CD27-TotalSeq-C | BioLegend | Clone O323 |
| Anti-CD21-TotalSeq-C | BioLegend | Clone Bu32 |
| Anti-IgD-TotalSeq-C | BioLegend | Clone IA6-2 |
| Anti-CD38-TotalSeq-C | BioLegend | Clone HIT2 |
| Anti-CD73-TotalSeq-C | BioLegend | Clone AD2 |
| Anti-CD85j-TotalSeq-C | BioLegend | Clone GHI/75 |
| Anti-PE Streptavidin-TotalSeq-C | BioLegend | CAT# 504155 |
| Anti-CD45RB-TotalSeq-C | BioLegend | Clone MEM-55 |
| Anti-CD10-TotalSeq-C | BioLegend | Clone HI10a |
| Anti-human Hashtag 1 Antibody | BioLegend | CAT#394661 |
| Anti-human Hashtag 2 Antibody | BioLegend | CAT#394663 |
| Anti-human Hashtag 3 Antibody | BioLegend | CAT#394665 |
| Anti-human Hashtag 4 Antibody | BioLegend | CAT#394667 |
| Anti-human Hashtag 5 Antibody | BioLegend | CAT#394669 |
| Anti-human Hashtag 6 Antibody | BioLegend | CAT#394671 |
| Anti-human Hashtag 7 Antibody | BioLegend | CAT#394673 |
| Anti-human Hashtag 8 Antibody | BioLegend | CAT#394675 |
| Anti-human Hashtag 9 Antibody | BioLegend | CAT#394677 |
| **Chemicals and Recombinant Proteins** |  |  |
| FcR Blocking Reagent | Miltenyi Biotec | CAT#130-059-901 |
| Rainbow Calibration Particles | ThermoFisher | CAT#A34305 |
| DRAQ7 Dead cell exclusion dye | BioLegend | CAT# 424001 |
| Streptavidin-APC conjugate | ThermoFisher | CAT#S32362 |
| Streptavidin-PE conjugate | BioLegend | CAT#405204 |
| Strep-Tactin APC | IBA-Lifesciences | CAT#6-5010-001 |
| Strep-Tactin PE | IBA-Lifesciences | CAT#6-5010-001 |
| **Experimental Models: Cell Lines** |  |  |
| Human Expi293F | Thermo Fisher | CAT#A14527 |
| Recombinant DNA |  |  |
| pVRC-SARS-CoV-2-RBD WA1/2020<br>and Omicron variant | Schmidt Lab | N/A |
| pCAGGS-SARS-CoV-2 Spike WA-1 2020<br>and Omicron variant | McClellan/Balazs/<br>Schmidt Labs | N/A |
| Software and Algorithms |  |  |
| FlowJo | FlowJo | www.flowjo.com |
| FACSDIVA<br>www.bdbiosciences.com/en-us/instruments/research-instruments/research-software/flow-cytometry-acquisition/facsdiva-software | BD Biosciences |  |
| Prism 8<br>www.graphpad.com/scientific-software/prism/ | GraphPad Software |  |
| CellRanger (v6.0.1)<br>https://support.10xgenomics.com/single-cell-vdj/software/ | 10X Genomics |  |
| R (v4.1.1) | R | www.r-project.org/ |
| Seurat (v4.0.6) | Seurat | satijalab.org/seurat/ |
| IgBlast (online server)<br>www.ncbi.nlm.nih.gov/igblast/ | National Library of Medicine |  |

### RESOURCE AVAILABILITY

#### Lead Contact

Further information and requests for resources and reagents should be directed to and will be fulfilled by the Lead Contact, Dr. Shiv Pillai

#### Materials Availability

The SARS-CoV-2 RBD and Spike proteins were made freely available through Dr. Aaron Schmidt during the pandemic but are now available from many commercial sources.

#### Data and Code Availability

The published article includes all data generated or analyzed during this study, and summarized in the accompanying tables, figures and Supplemental materials.

Sequence data was deposited in the GEO database.

The accession number for sequence information is GSE 278378.

No unique codes were generated during analysis of the data.

## Methods

### Human Samples

Peripheral blood samples were drawn from patients with COVID-19 at Massachusetts General Hospital and vaccinees at MGH and the University of Massachusetts Medical Center. PBMCs were stored in liquid nitrogen for subsequent batch experiments. For all vaccination cohorts, prior exposure to SARS-CoV-2 was determined by measuring serologic responses to SARS-CoV-2 Spike and Nucleocapsid antigens in baseline, pre-vaccination plasma samples. Antibodies to the spike protein receptor binding domain and nucleocapsid were measured by the Roche Elecsys Anti-SARS-CoV-2 S and Anti-N assay (Roche Diagnostics, Indianapolis, IN), per manufacturer’s instructions. Uninfected and unvaccinated subjects were defined by having no detectable anti-spike or anti-nucleocapsid antibodies. Acute COVID-19 was defined by active or resolving symptoms of pneumonia and ongoing hospitalization for SARS-CoV-2 infection as confirmed by RT-PCR. The median number of days from symptom onset to blood draw was 28 and the interquartile range was 14 to 29 days. Convalescence was defined as a clinically asymptomatic state on the date of blood draw, either from a baseline asymptomatic state or recuperated from clinical symptoms of COVID-19. Fifty eight of 66 convalescent donors had symptomatic infection with well-defined dates of symptom onset. For these donors, the median number of days from symptom onset to blood draw was 217 and the interquartile range was 174 to 248 days. 8 of the 66 convalescent COVID-19 patients had been previously asymptomatic but were identified by the presence of anti-SARS-CoV-2 spike or anti-nucleocapsid responses at the baseline, and duration of infection could not be determined. Demographic and clinical data related to vaccinated and COVID-19 cohorts are detailed in **Tables S1-S2**. Healthy donor samples used in Figures 2C-D were all historic PBMCs collected prior to March of 2020. Healthy donor was defined by the lack of any ongoing or historic infectious, autoimmune, or malignant condition, as determined by review of the medical record. This cohort (n=21) had a median age of 66 (IQR 60-74) and consisted of 52% males and 48% females.

Sample pairing: Among vaccinee cohorts, 47% of week 3-4 post dose 1 samples were paired with subjects from week 1-2 post dose 1, 89% of week 1-2 post dose 2 samples were paired with subjects from week 3-4 post dose 1, and 92% of week 4-8 post dose 2 samples were paired with subjects from week 1-2 post dose 2. Five of 6 subjects from the week 1 post dose 3 time point were paired with subjects from the week 24 post dose 2 time point. Additional details related to sample pairing across cohorts are available in **Table S1**.

### Study Approval

This study was performed with the approval of the Institutional Review Boards at the Massachusetts General Hospital and the University of Massachusetts Medical School.

### Protein Expression and Purification

SARS-2 RBD (GenBank: MN975262.1) was cloned into pVRC vector containing an HRV 3C-cleavable C-terminal His8X and Avi-tags and sequence confirmed by Genwiz. SARS-CoV-2 full-length spike protein contained a C-terminal foldon trimerization domain and His8X and Avi tags (Wrapp et al., 2020). The constructs were transiently transfected into mammalian Expi293F suspension cells for recombinant expression. Five to seven days post-transfection, supernatants were harvested by centrifugation and purified by immobilized metal affinity chromatography (IMAC) using Cobalt-TALON resin (Takara) followed by size exclusion chromatography on Superdex 200 Increase 10/300 GL (GE Healthcare) in PBS. Purity was assessed by SDS-PAGE analysis. Purified RBD proteins were site-specifically biotinylated using the BirA biotin-protein ligase kit (Avidity) following manufacturer’s instructions.

### Flow cytometry and Tetramer Generation

Cryopreserved PBMCs in aliquots of 5-10 million cells were thawed quickly at 37℃ and DMSO rapidly diluted with 1% BSA in PBS. Cells were incubated at 21℃ for 15 minutes prior to transfer to FACS tubes for centrifugation. All centrifugation steps were carried out at 1,250 RPM at 4 ℃. Prior to antibody staining, Fc receptors were blocked using human FcR blocking reagent (Miltenyi) at a concentration of 1:20 at 4℃ for 10 minutes. For antigen-specific experiments, we generated two distinct antigen-specific tetramer probes, each labeled with a different fluorophore, combined with a broader flow cytometry panel to characterize all class-switched B cell subsets. For all spike-specific studies, tetramers were generated using Strep-Tagged proteins and fluorescent Strep-Tactin probes. Strep-Tactin APC (IBA-Lifesciences, CAT#6-5010-001) and Strep-Tactin PE (IBA-Lifesciences, CAT#6-5010-001) were used for dual labeling of antigen-specific B cells. An optimal staining ratio of Strep-Tagged antigen to Strep-Tactin probe of 25 nM antigen to 0.5 uL Strep-Tactin was empirically determined and used consistently for all experiments.

For RBD-specific studies, tetramers were generated using site-specific biotinylated proteins and fluorescent streptavidin probes. Wild-type RBD was separately complexed with Streptavidin-APC (ThermoFisher, CAT#S32362) and Streptavidin-PE (BioLegend, CAT#405204). Complexes were formed using an antigen to streptavidin molar ratio of 4:1 for all RBD experiments.

Tetramers were generated at the time of each experiment. Each antigen plus probe dilution was mixed as described above and incubated at 4℃ for 45-minutes protected from light. Cells were surface stained first at 37℃, protected from light, using optimized concentrations of fluorochrome-conjugated primary antibodies for 20 minutes. Cells were then washed, centrifuged, and resuspended in a separate 4℃ antibody master mix for an additional 30-minute stain at 4℃. Antigen complexes were added to the 4℃ master mix. Finally, cells were washed, centrifuged, and re-suspended in DRAQ7 dead cell exclusion dye (BioLegend) at a concentration of 1:1000. DRAQ7 is a membrane-impermeable dye with an emission maxima of 678/697 nm once intercalated with dsDNA. We designed our flow panel with CD3 on APC/R700 and CD19 on APC/Cy7 such that any cells with dual-emission across these two channels was considered non-viable and excluded (as displayed in **Figure S1**). As shown in the comprehensive gating strategy **(Figure S1)**, CD38 high plasmablasts were not quantitated during flow cytometric analyses because our primary focus was to identify memory B cell subsets and the staining approach employed was not designed for optimal capture of plasmablasts with low or no surface BCR. Antibodies used with manufacturer, clone, conjugate and staining information are as follows: anti-human CD3 (Miltenyi Biotec, Clone BW264/56, APC-R700, 1:200, 4℃); anti-human CD19 (BioLegend, Clone SJ25C1, APC-Cy7, 1:25, 4℃); anti-human CD27 (BD Biosciences, Clone L128, BUV395, 1:40, 4℃); anti-human CD27 (BioLegend, Clone M-T271, PE-Dazzle, 1:100, 4℃); anti-human IgD (BioLegend, Clone IA6-2, BV785, 1:100, 37℃); anti-human-CD38 (BioLegend, Clone HIT2, BV421, 1:100, 4℃); anti-human-CD21 (BD Biosciences, Clone B-ly4, BV480, 1:100, 37℃); anti-human CD11c (BD Biosciences, Clone B-ly6, BV605, 1:100, 4℃); anti-human CXCR5 (BD Biosciences, Clone RF8B2, A488, 1:100, 37℃), anti-human CD20 (BD Biosciences, Clone 2H7, BUV805, 1:100, 37℃); anti-human IgG (BD Biosciences, Clone G18-145, BB700, 1:100, 4℃); anti-human IgA (Miltenyi Biotec, Clone IS11-8E10, APC, 1:100, 4℃); anti-human IgM (BD Biosciences; Clone G20-127, BUV395, 1:25, 4℃). All flow cytometry was performed on a BD Symphony cytometer (BD Biosciences, San Jose, CA) and rainbow tracking beads (8 peaks calibration beads, Fisher) were used to ensure consistent cytometer signal output between flow cytometry experiments.

### Bulk RNA Sequencing

Naïve (CD19+ CD20+ IgD+ CD27-CXCR5+ CD11c-), DN1 (CD19+ CD20+ IgD- CD27- CXCR5+ CD11c-), and SWM (CD19+ CD20+ IgD-CD27+ CXCR5+ CD11c-) B cells were FACS purified from cryopreserved PBMC’s using a FACS Aria Fusion cell sorter. Total RNA was isolated from sorted B cell populations using the RNAeasy Plus Micro kit (Qiagen) according to the manufacturer’s instructions. RNA quality was assessed via Agilent 4200 Tapestation and RNA concentration was determined using a Qubit Flex (Fisher Scientific). Total RNA (4 ng/sample) was processed using the SMARTer Stranded Total RNA-seq Kit v3-Pico Input Mammalian (Takara). Final libraries were quantified with a Tapestation (Agilent) prior to dilution and pooling for sequencing on a NextSeq 2000 (Illumina) using 2x100 bp paired-end chemistry.

### RNA sequencing analysis

The reads in each sample were mapped to human reference genome GrCh38 using STAR software (version 2.7.9a) [Dobin, 2013]. On average, we obtained ∼11.3 million uniquely mapped reads per sample. Raw read count matrix (the number of reads for each gene in each sample) was obtained using HTSeq python package (version 2.0.1) [Putri, 2022]. The genes with less than 5 reads on average in all samples were removed, resulting 26,777 genes remaining for further analysis. The differential expression analyses between different cell populations were performed using likelihood ratio test (LRT) in DESeq2 R package (version 1.38.1). Genes with FDR-adjusted p-value < 0.05 were considered differentially expressed genes (DEG). Raw read count data was normalized using variance stabilizing transformation (VST) to obtain VST-transformed data (VSD), which was used for generating principal component analysis plots and visualization of individual gene expression. Pathway enrichment analysis was performed using ReactomePA R package (version 1.41.1) [Yu and He, 2016]. The regulator enrichment analysis was performed using RegEnrich R package (version 1.8.0) [Tao, 2022].

### Bulk BCR Sequencing

Naïve (CD19+ CD20+ IgD+ CD27-CXCR5+ CD11c-), DN1 (CD19+ CD20+ IgD-CD27-CXCR5+ CD11c-), and SWM (CD19+ CD20+ IgD-CD27+ CXCR5+ CD11c-) B cells were FACS purified from freshly isolated PBMC’s using a FACS Aria Fusion cell sorter. Cells were directly sorted into DNA LoBind tubes containing DNA RNA Sheild (Zymo). Genomic DNA was isolated from each lysate using Quick-DNA/RNA Microprep Plus Kits (Zymo), per manufacturer’s protocol. Full-length heavy chain sequences were amplified using equimolar VH and JH primer cocktails and Multiplex PCR Kits (Qiagen), as previously described (Rosenfeld et al., 2022). IGH PCR products were enriched using AMPure bead purification following the manufacturer’s protocol (Beckman Coulter). Illumina barcodes and indices were added with a 2nd round of PCR using Multiplex PCR Kits (Qiagen), as previously described (Rosenfeld et al., 2022). Barcoded PCR products were run through 2% agarose gel and gel purified using a QIAquick Gel Extraction Kit (Qiagen). Each PCR product was quantified on Qubit using a dsDNA high-sensitivity kit and equimolar libraries were prepared. Libraries were sequenced using a 600 cycle MiSeq Kit v3 and an Illumina MiSeq sequencer.

For Antibody gene repertoire analyses, IGHV gene BCR sequences were aligned with MiXCR (Bolotin et al., 2015) and subsequently analyzed with VDJtools (Shugay et al., 2015) and Shazam, a part of the Immcantation framework for Adaptive Immune Receptor Repertoire (AIRR) analyses (Gupta et al. 2015).

### TP63 Quantitative PCR

From the same cell lysates described under the bulk BCR sequencing section, RNA was purified using Quick-DNA/RNA Microprep Plus Kits (Zymo), per manufacturer’s protocol. Reverse transcription was performed using Applied Biosystems’ High-Capacity cDNA Reverse Transcriptino kit (catalogue #4374966). Real-time PCR was next performed using Applied Biosystems’ TaqMan Gene Expression Assay for TP63 (Hs00978340_m1) and TaqMan Fast Advanced Master Mix (catalogue #4444557). Thermocycling was carried out using a BioRad T100 Thermal Cycler for 40 cycles and Applied Biosystems’ ViiA 7 Real-Time PCR System was used to analyze the expression of TP63.

### Quantification and Statistical Analyses

FCS files were analyzed, and B cell subsets were quantified using FlowJo software (version 10). GraphPad Prism version 8 was used for all statistical analyses and plotting of data points. For comparison of paired samples, the Wilcoxon matched-pairs signed rank test, or the Friedman test was used to calculate p-values, depending on the distribution of the data. For comparing unpaired data sets, the Mann Whitney U test was used. A p-value of < 0.05 was considered significant.

## Supplemental Information Titles and Legends

**Supplemental Figure 1: Flow cytometry hierarchical gating strategy.** Pseudocolor and contour plots displaying hierarchical gating strategy used to quantify B cell subsets. For each sample, gating was defined using total B cells and then applied to antigen-specific B cells.

**Supplemental Figure 2: SARS-CoV-2 RBD-specific B cells are absent while CD27^+^ SWM and DN1 B cells dominate spike-specific B cells among SARS-CoV-2 naïve hosts.** A) Representative pseudocolor plots of RBD-specific B cells among unvaccinated, SARS-CoV-2 naïve hosts (N=6), mRNA vaccine recipients post-boost (N=3), and convalescent COVID-19 patients (N=3) demonstrating the absence of RBD-specific B cells among SARS-CoV-2 naïve hosts. B) Representative pseudocolor plots of spike-specific B cells among unvaccinated, SARS-CoV-2 naïve hosts (N=6). C) Donut charts displaying the proportion of CD27^+^ SWM and DN1 B cells among total spike-specific B cells from unvaccinated, SARS-CoV-2 naïve hosts (N=6). Each donut chart represents the B cell profile from one individual donor. Numbers provided within each donut chart represent the total number of spike-specific B cells profiled for each donor.

**Supplemental Figure 3: DN1 B cells are distinguished from canonical CD27^+^ SWM by higher expression of CD62L and CXCR4 with lower expression of IGF-1R.** A) Mean fluorescent intensity (MFI) data for CXCR5, CD11c, CD86, CD20, CD27, CD40, CD62L, CXCR4, and IGF-1R across B cell subsets. B) Normalized gene expression data for *SELL, CXCR4,* and *IGF1R* from bulk RNA sequencing of B cell subsets.

**Supplemental Figure 4: Preferential usage of different metabolic pathways may distinguish memory B cell subsets.** Dot plot displaying gene set enrichment analysis comparing DN1 B cells to CD27^+^ SWM and naïve B cells. Size of dot indicates number of genes enriched within respective gene set. A) Dot plot displaying gene sets over-expressed by DN1 B cells. B) Dot plot displaying gene sets under-expressed by DN1 B cells. C) Normalized gene expression data for *TP53* and *TP73* from bulk RNA sequencing of B cell subsets.

**Supplemental Figure 5: Memory B cell subsets may differ in their predilection for class switching to IgA and IgG2**. Normalized gene expression data for *IGHG1, IGHG2, IGHG3, IGHG4, IGHA1, IGHA2,* and *IGHE* from bulk RNA sequencing of B cell subsets.

**Supplemental Table 1: Vaccinee Cohort Demographic Data**

**Supplemental Table 2: COVID-19 Cohort Demographic Data**

