## Supplemental Figures and Tables for "Two distinct durable human class-switched memory B cell populations are induced by vaccination and infection"

**Supplemental Figure 1**

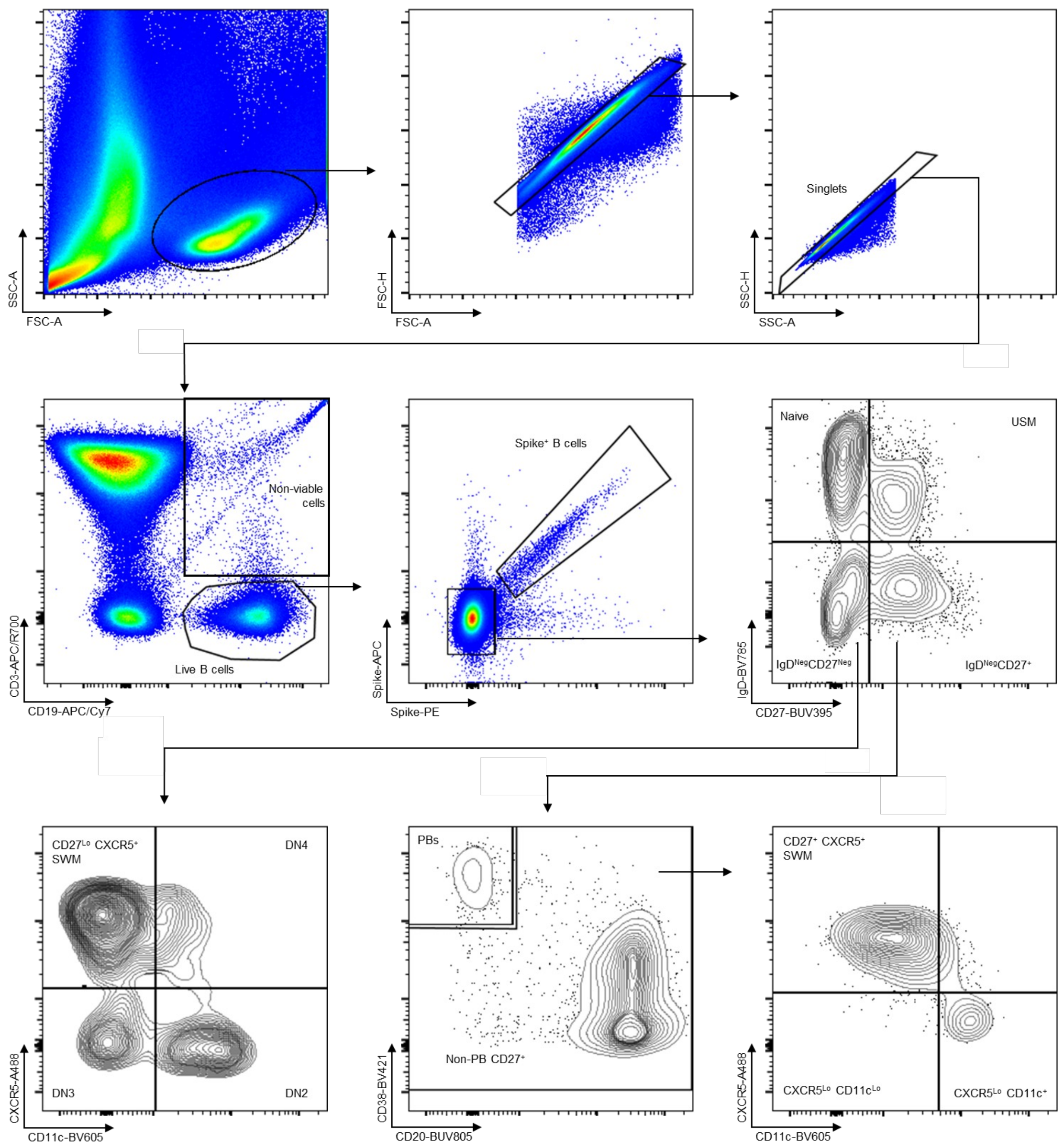


**Flow cytometry hierarchical gating strategy**. Pseudocolor and contour plots displaying hierarchical gating strategy used to quantify B cell subsets. For each sample, gating was defined using total B cells and then applied to antigen-specific B cells.

**Supplemental Fig 2**

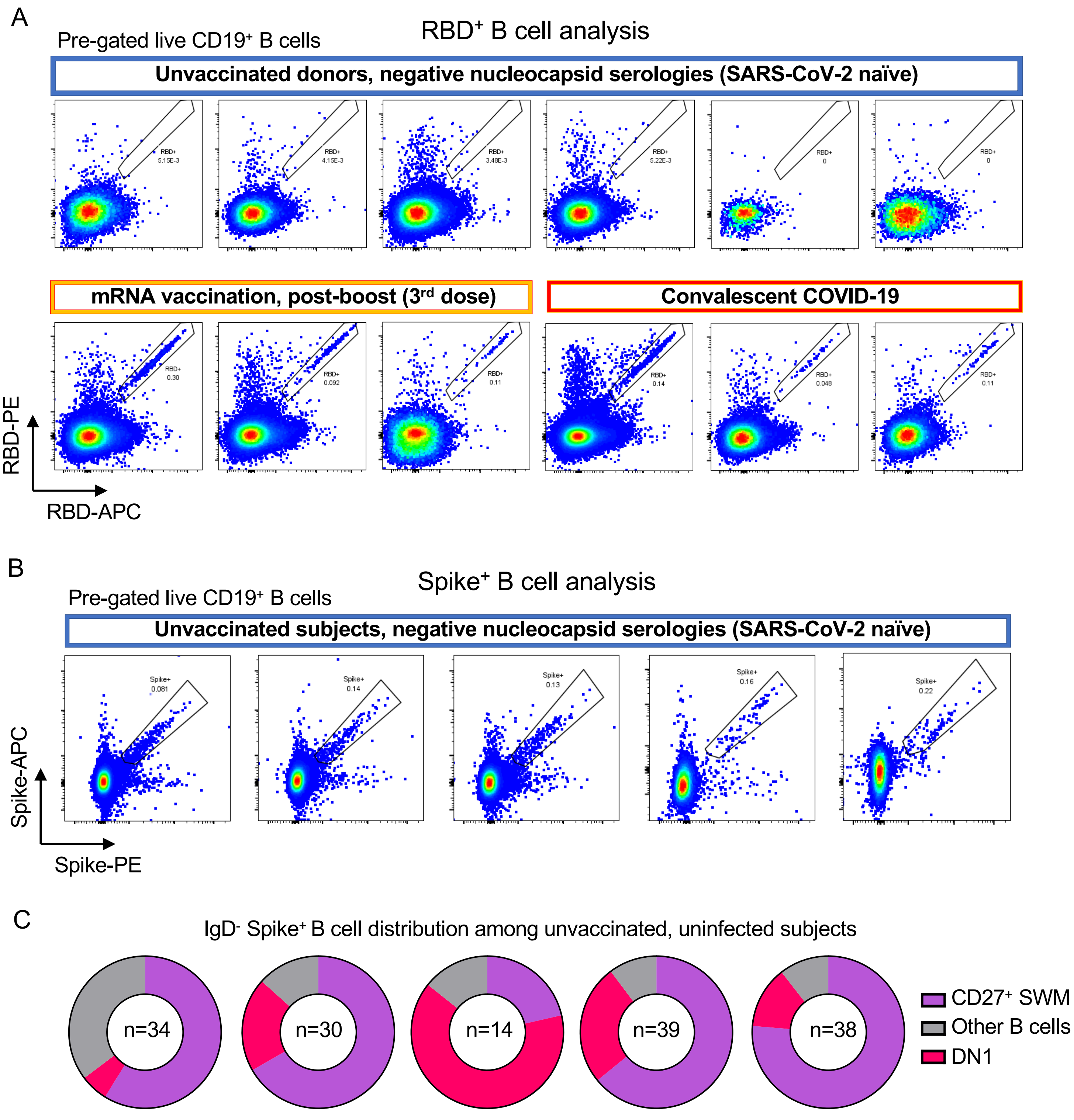


**SARS-CoV-2 RBD-specific B cells are absent while CD27+ SWM and DN1 B cells dominate spike-specific B cells among SARS-CoV-2 naïve hosts.** A) Representative pseudocolor plots of RBD-specific B cells among unvaccinated, SARS-CoV-2 naïve hosts (N=6), mRNA vaccine recipients post-boost (N=3), and convalescent COVID-19 patients (N=3) demonstrating the absence of RBD-specific B cells among SARS-CoV-2 naïve hosts. B) Representative pseudocolor plots of spike-specific B cells among unvaccinated, SARS-CoV-2 naïve hosts (N=6). C) Donut charts displaying the proportion of CD27+ SWM and DN1 B cells among total spike-specific B cells from unvaccinated, SARS-CoV-2 naïve hosts (N=6). Each donut chart represents the B cell profile from one individual donor. Numbers provided within each donut chart represent the total number of spike-specific B cells profiled for each donor.

**Supplemental Figure 3**

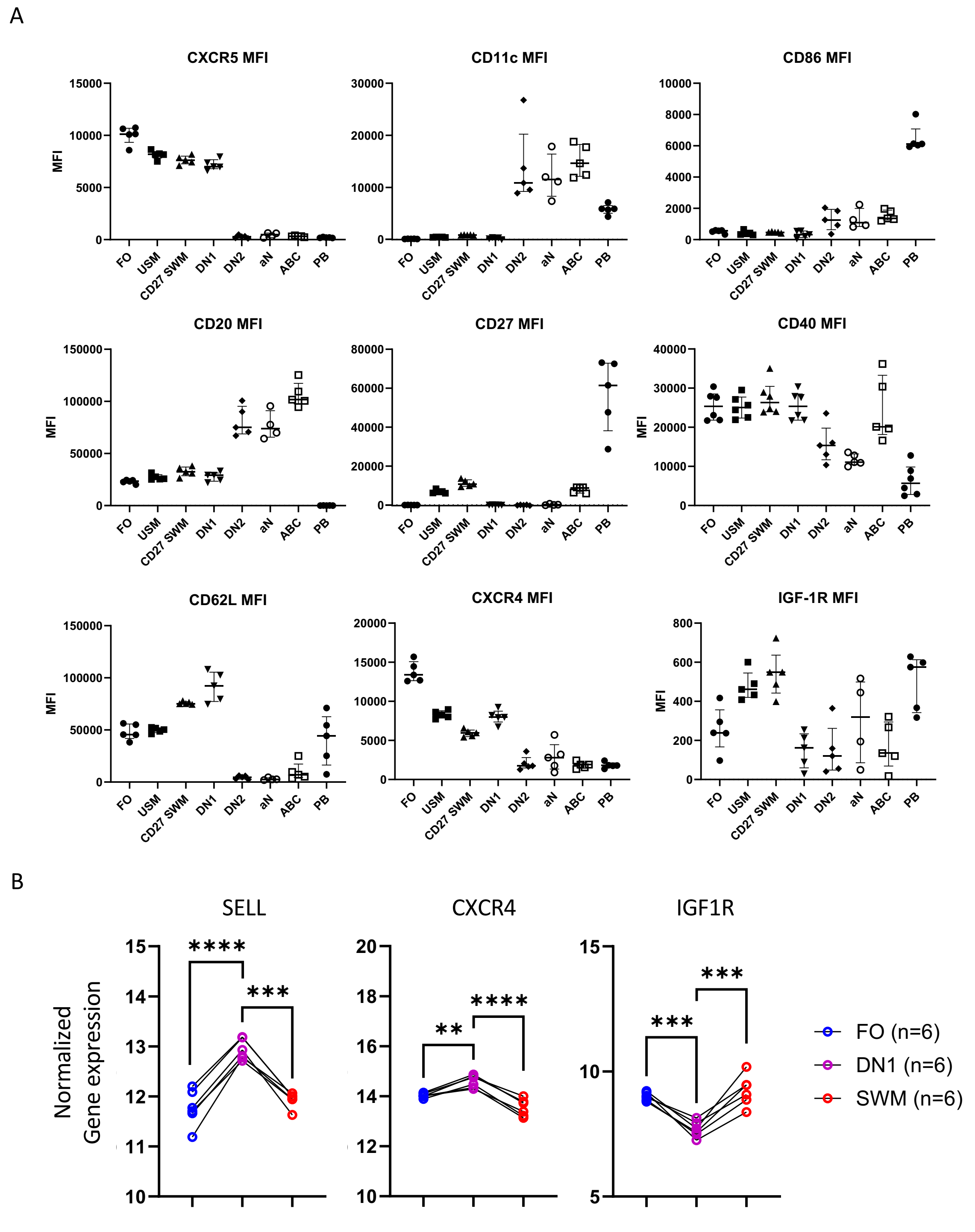


**DN1 B cells are distinguished from canonical CD27+ SWM by higher expression of CD62L and CXCR4 with lower expression of IGF-1R.** A) Mean fluorescent intensity (MFI) data for CXCR5, CD11c, CD86, CD20, CD27, CD40, CD62L, CXCR4, and IGF-1R across B cell subsets. B) Normalized gene expression data for SELL, CXCR4, and IGF1R from bulk RNA sequencing of B cell subsets.

**Supplemental Figure 4**

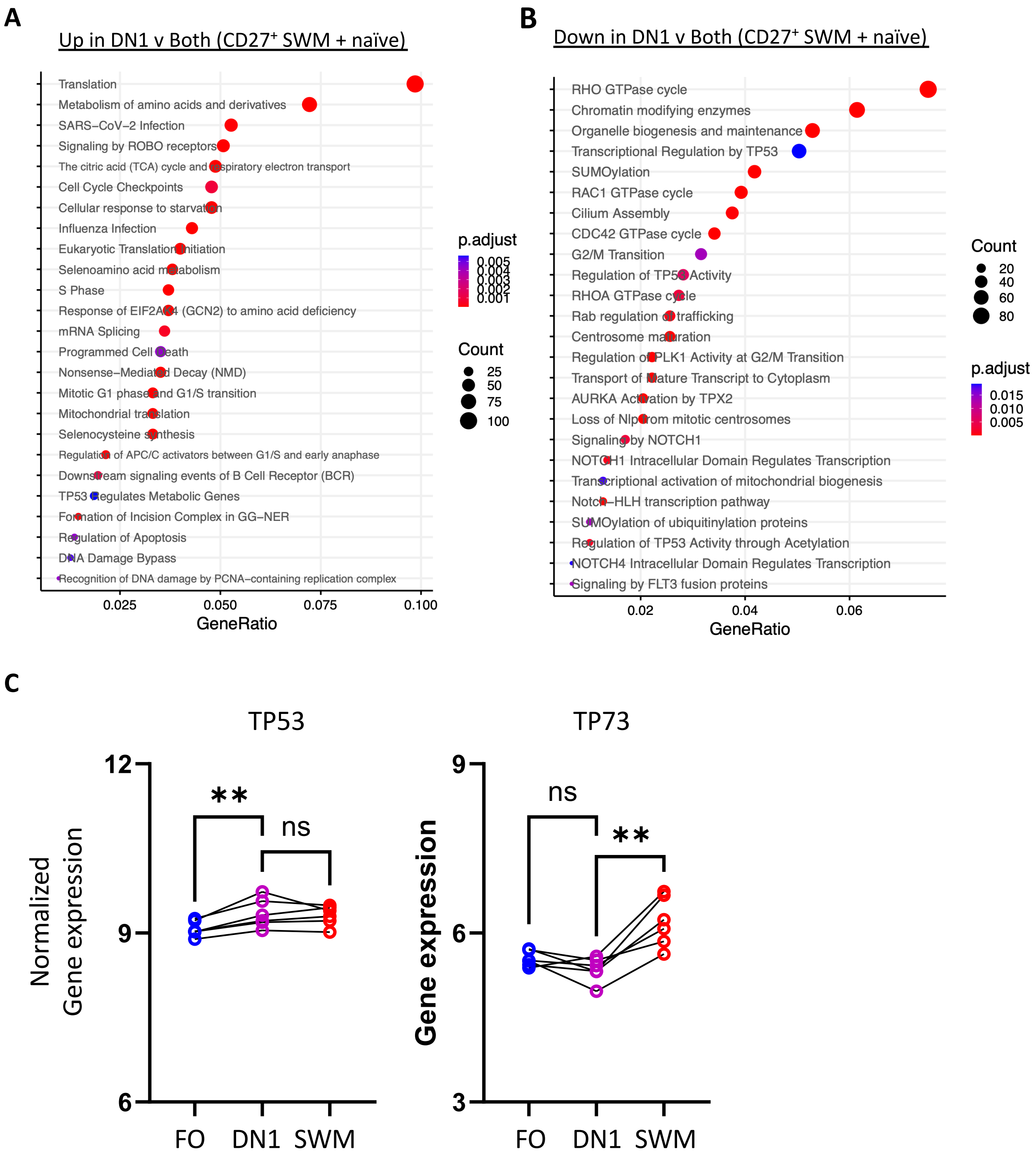


**Preferential usage of different metabolic pathways may distinguish memory B cell subsets.** Dot plot displaying gene set enrichment analysis comparing DN1 B cells to CD27+ SWM and naïve B cells. Size of dot indicates number of genes enriched within respective gene set. A) Dot plot displaying gene sets over-expressed by DN1 B cells. B) Dot plot displaying gene sets under-expressed by DN1 B cells. C) Normalized gene expression data for *TP53* and *TP73* from bulk RNA sequencing of B cell subsets.

**Supplemental Figure 5**

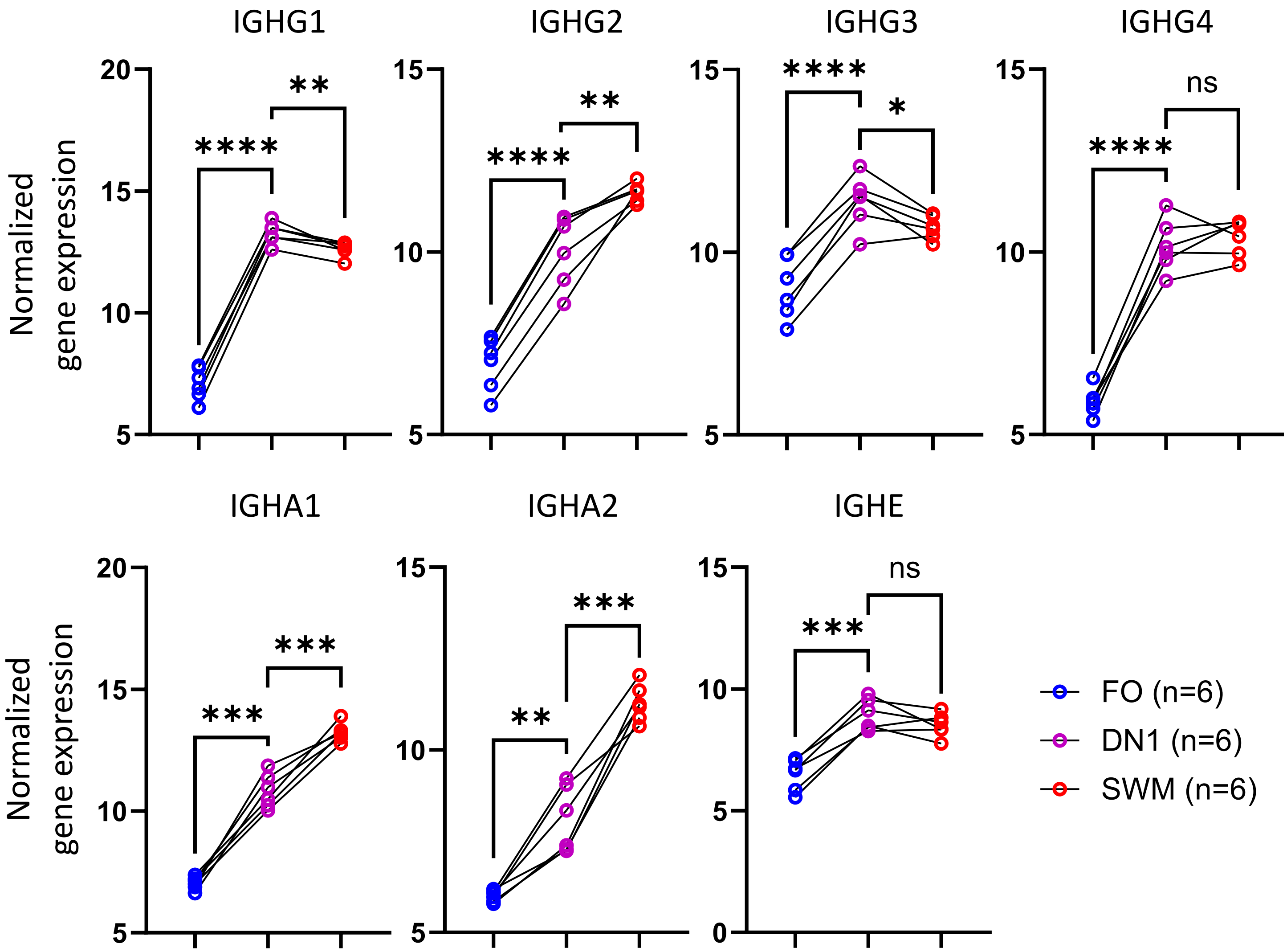


**Memory B cell subsets may differ in their predilection for class switching to IgA and IgG2**. Normalized gene expression data for *IGHG1, IGHG2, IGHG3, IGHG4, IGHA1, IGHA2,* and *IGHE* from bulk RNA sequencing of B cell subsets.

**Supplemental Table 1**

**Vaccinee Cohort Demographic Data**

| Vaccinee Cohort Demographic Data | | |
|---|---|---|
| Pre-vaccination baseline (n=11) | | |
| | Male, n (%) | 3 (27) |
| | Female, n (%) | 8 (73) |
| | Caucasian, n (%) | 6 (55) |
| | African American, n (%) | 2 (18) |
| | Hispanic, n (%) | 3 (27) |
| | Age, median (IQR) | 51 (31-63) |
| | Prior SARS-CoV-2 infection, n (%) | 0 (0%) |
| Week 1-2 post-dose 1 (n=9) | | |
| | Male, n (%) | 5 (56) |
| | Female, n (%) | 4 (44) |
| | Caucasian, n (%) | 8 (89) |
| | African American, n (%) | 1 (11) |
| | Hispanic, n (%) | 0 (0) |
| | Age, median (IQR) | 56 (43-64) |
| | Prior SARS-CoV-2 infection, n (%) | 0 (0) |
| | Days from last immunization, median (IQR) | 9 (7.5-10) |
| | Paired samples from baseline, n (%) | 1 (11) |
| Week 3-4 post-dose 1 (n=19) | | |
| | Male, n (%) | 9 (47) |
| | Female, n (%) | 10 (53) |
| | Caucasian, n (%) | 17 (89) |
| | African American, n (%) | 2 (11) |
| | Hispanic, n (%) | 0 (0) |
| | Age, median (IQR) | 60 (45-71) |
| | Prior SARS-CoV-2 infection, n (%) | 0 (0) |
| | Days from last immunization, median (IQR) | 21 (20-27) |
| | Paired samples from baseline, n (%) | 5 (26) |
| | Paired samples from Week 1-2 post-dose 1, n (%) | 9 (47) |
| Week 1-2 post-dose 2 (n=19) | | |
| | Male, n (%) | 8 (42) |
| | Female, n (%) | 11 (58) |
| | Caucasian, n (%) | 17 (89) |
| | African American, n (%) | 2 (11) |
| | Hispanic, n (%) | 0 (0) |
| | Age, median (IQR) | 61 (52-74) |
| | Prior SARS-CoV-2 infection, n (%) | 0 (0) |
| | Days from last immunization, median (IQR) | 29 (28-35) |
| | Paired samples from baseline, n (%) | 5 (26) |
| | Paired samples from Week 1-2 post-dose 1, n (%) | 9 (47) |
| | Paired samples from Week 3-4 post-dose 1, n (%) | 17 (89) |
| Week 4-8 post-dose 2 (n=12) | | |
| | Male, n (%) | 7 (58) |
| | Female, n (%) | 5 (42) |
| | Caucasian, n (%) | 12 (100) |
| | African American, n (%) | 0 (0) |
| | Hispanic, n (%) | 0 (0) |
| | Age, median (IQR) | 66 (62-71) |
| | Prior SARS-CoV-2 infection, n (%) | 0 (0) |
| | Days from last immunization, median (IQR) | 62 (58-87) |
| | Paired samples from baseline, n (%) | 1 (8) |
| | Paired samples from Week 1-2 post-dose 1, n (%) | 7 (58) |
| | Paired samples from Week 3-4 post-dose 1, n (%) | 11 (92) |
| | Paired samples from Week 1-2 post-dose 2, n (%) | 11 (92) |
| Week 24 post-dose 2 (n=10) | | |
| | Male, n (%) | 3 (30) |
| | Female, n (%) | 7 (70) |
| | Caucasian, n (%) | 8 (80) |
| | African American, n (%) | 0 (0) |
| | Hispanic, n (%) | 2 (20) |
| | Age, median (IQR) | 44 (38-65) |
| | Prior SARS-CoV-2 infection, n (%) | 0 (0) |
| | Days from last immunization, median (IQR) | 223 (180-287) |
| | Paired samples from baseline, n (%) | 0 (0) |
| | Paired samples from Week 1-2 post-dose 1, n (%) | 0 (0) |
| | Paired samples from Week 3-4 post-dose 1, n (%) | 0 (0) |
| | Paired samples from Week 1-2 post-dose 2, n (%) | 0 (0) |
| | Paired samples from Week 4-8 post-dose 2, n (%) | 0 (0) |

**Supplemental Table 2**

### COVID-19 Cohort Demographic Data

| COVID-19 Cohort Demographic Data | | |
|---|---|---|
| Acute COVID-19 (n=13) | | |
| | Male, n (%) | 7 (54) |
| | Female, n (%) | 6 (46) |
| | Caucasian, n (%) | 7 (54) |
| | African American, n (%) | 1 (8) |
| | Hispanic, n (%) | 7 (54) |
| | Age, median (IQR) | 62 (45-73) |
| | Outpatient symptomatic infection, n (%) | 0 (0) |
| | Hospital admission, regular floor, n (%) | 0 (0) |
| | Hospital admission, intensive care, n (%) | 13 (100) |
| | Vaccinated, n (%) | 0 (0) |
| | Days from symptom onset, median (IQR) | 23 (15-29) |
| Convalescent COVID-19 (n=66) | | |
| | Male, n (%) | 24 (36) |
| | Female, n (%) | 42 (64) |
| | Caucasian, n (%) | 48 (72) |
| | African American, n (%) | 1 (2) |
| | Hispanic, n (%) | 16 (24) |
| | Age, median (IQR) | 39 (32-56) |
| | Asymptomatic infection, n (%) | 8 (12) |
| | Outpatient symptomatic infection, n (%) | 46 (70) |
| | Hospital admission, regular floor, n (%) | 7 (11) |
| | Hospital admission, intensive care, n (%) | 5 (8) |
| | Vaccinated, n (%) | 0 (0) |
| | Days from symptom onset, median (IQR) | 217 (174-248) |
